# MeCP2 binding and genome–lamina reorganization precede long gene activation during mouse corticogenesis

**DOI:** 10.1101/2025.09.01.673434

**Authors:** Pim M. J. Rullens, Sandra S. de Vries, Silke J. A. Lochs, Corina M. Markodimitraki, Keith M. Garner, Youri Adolfs, Hegias Mira-Bontenbal, Micha M. Müller, Elisa Bugani, Marvin E. Tanenbaum, Joost Gribnau, Onur Basak, Jop Kind

## Abstract

During corticogenesis, neural gene expression is tightly coordinated by chromatin and epigenetic changes, whose misregulation can lead to neurodevelopmental disorders^1–4^. The role of spatial genome organization—particularly interactions with the nuclear lamina—during these developmental programs remains poorly understood. Here, we combined in utero electroporation with scDam&T-seq to jointly profile genome–lamina contacts and transcriptomes in single cells of the mouse embryonic cortex. Interestingly, we find extensive genome–lamina reorganization during corticogenesis that is strongly biased towards long neuronal genes (≥100 kb), which are associated with neurodevelopmental disorders including autism spectrum disorder. Detachment of these genes frequently precedes transcriptional activation, positioning lamina disengagement as an early gene regulatory event. We identify the methyl CpG binding protein 2 (MeCP2)—mutated in Rett syndrome—as a candidate mediator of this process. MeCP2 binds lamina-associated, hydroxymethylated long genes before their repositioning, suggesting that MeCP2 may play a role in genome–lamina reorganization. These findings suggest a link between prevalent genome–lamina reorganization and MeCP2 regulation to ensure proper spatiotemporal activation of long neuronal genes during corticogenesis.

## Main

The formation of the cerebral cortex, or corticogenesis, is a complex and tightly regulated developmental process that establishes the functional architecture of the mammalian brain. During this period, neural progenitor cells undergo changes in chromatin organization, gene expression, and cellular architecture as they differentiate into specialized cortical cell types^1–4^. In recent years, significant advances have increased our understanding of how chromatin organization and epigenetic regulation influence gene regulation during corticogenesis^2,5,6^. However, the role of spatial genome organization in this process remains poorly understood.

Previous studies suggest that the spatial compartmentalization of the genome within various nuclear bodies plays a critical role in the gene regulatory programs that drive development^7–11^. Of particular interest is the interaction between the genome and the nuclear lamina—a filamentous protein meshwork lining the inner nuclear membrane—which has emerged as a key structural component of chromatin architecture and a potential regulator of transcriptional activity^12–15^. However, major gaps remain in our understanding of how genome–lamina contacts contribute to the dynamic gene regulation that is essential for brain development. Gaining insight into how genome–lamina contacts shape gene expression programs during corticogenesis may illuminate mechanisms underlying both normal brain development and the etiology of neurodevelopmental disease. Addressing this challenge requires experimental approaches capable of simultaneously profiling genome–lamina contacts and transcription at single-cell resolution.

In this study, we jointly profiled genome–lamina contacts and transcriptomes in single cells from the developing mouse cerebral cortex during neurogenesis, a critical period marked by extensive transcriptional changes. We find a large cohort of long, neuronal function-related genes that undergo spatial genome–lamina repositioning during neurogenesis. Importantly, we observe that detachment of these genes often precedes their transcriptional activation, suggesting that release from the lamina is an early regulatory step in long neuronal gene activation. Furthermore, our analyses link genome–lamina reorganization to the transcriptional regulator MeCP2 – a gene mutated in the neurodevelopmental disorder Rett syndrome^16^ – and its methylated DNA ligands. Single-cell profiling of MeCP2 in the developing mouse cortex, revealed that MeCP2 binds preferentially to lamina–associated, hydroxymethylated long genes prior to their release from the lamina and subsequent transcriptional activation.

Together, our findings reveal genome–lamina repositioning as a prevalent and early layer of gene regulation during cortical development—one that is potentially influenced by MeCP2 and disrupted in the context of Rett syndrome.

### In utero electroporation scDam&T-seq enables joint profiling of LADs and transcription in single cells of the mouse embryonic cortex

To study the role of genome–lamina contacts, or Lamina-associated domains (LADs), during cortical development, we designed an approach that employs in utero electroporation (IUE) to perform scDam&T-seq^17,18^ in vivo in the mouse embryonic cortex (Fig. 1a). To this end, we used a DamID construct that fuses Dam to the core nuclear lamina component Lamin B1^13^. In addition, we aimed to dissect the interplay between LADs and chromatin accessibility through IUE and scDam&T-seq with a DamID construct encoding the Dam enzyme alone. The untethered Dam enzyme has previously been reported to specifically mark accessible DNA, allowing for the accurate profiling of open chromatin regions^18,19^. For this purpose, we employed a catalytically impaired variant of Dam, mutant N126A (henceforward Dam126), that increases signal specificity^20^. We performed IUE on the ventricular zone (VZ) of the embryonic cortex throughout neurogenesis, at embryonic day (E) 13, 14 and 16. Apical progenitors (AP) reside in the VZ and play a crucial role during neurogenesis, as they are highly proliferative and have the potential to self-renew and differentiate into intermediate progenitors and neurons^21^. By predominantly targeting APs with IUE at these embryonic stages, we aimed to express the DamID constructs throughout both multipotent and differentiating cortical cell types. After IUE, the embryos were left to develop for another 48 hours before the collection and processing of the cortical tissue. First, we examined the nuclear expression of the Dam-Lamin B1 fusion protein in cortical tissue slices, using immunofluorescence imaging. As expected, Dam-Lamin B1 is exclusively expressed at the nuclear periphery, corroborating proper incorporation into the lamina meshwork (Fig. 1b). Importantly, expression of Dam-Lamin B1 is visible across the cortical layers, from VZ up to the cortical plate (CP) (Fig. 1b).

**Fig 1.**
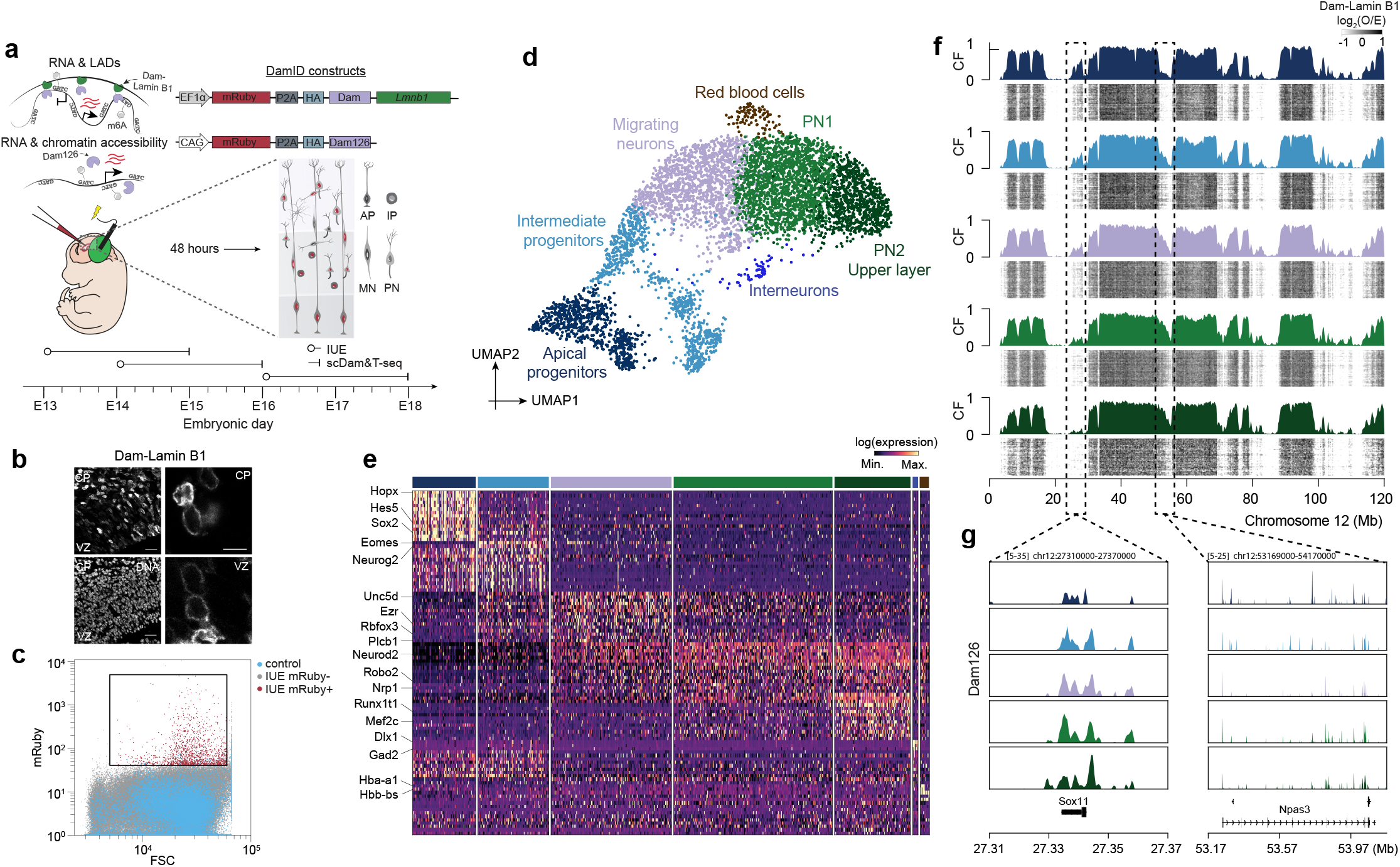
Joint profiling of genome–lamina contacts and transcriptomes in the developing mouse cortex. **a**, Overview of experimental design. DamID constructs are injected and electroporated in utero (IUE) into the ventricular zone of the mouse cerebral cortex at embryonic day (E) 13, 14 and 16. After 48 hours, samples are dissociated for scDam&T-seq analysis. Cartoon partially adapted from ref. 78. **b**, Immunofluorescence on HA-tag in the Dam-Lamin B1 DamID construct on cortical tissue slides at E16. Bottom left image shows a DAPI DNA staining. Left images are 20X magnifications and white scale bar is 25 µm, and right are 60X magnifications and white scale bar is 5 µm. **c**, FACS analysis on dissociated cortical tissue after IUE with Dam-Lamin B1. mRuby+ cells were sorted for scDam&T-seq (Methods). **d**, UMAP dimensionality reduction plot of identified cell types. **e**, Heatmap visualization showing RNA expression of top 15 marker genes of each cell type. **f**, Single-cell heatmaps of Dam-Lamin B1 data for different cell types along chromosome 12. Above each heatmap a contact frequency (CF) track. **g**, Chromosome 12 zoomins, showing cell type-specific pseudobulks of Dam126 data of the depicted loci.

Next, we dissociated cortical tissue to perform scDam&T-seq on FACS sorted mRuby+ cells (Fig. 1c, Extended Data Fig. 1a)(Methods). We obtained high quality multimodal data, spanning multiple biological replicates and embryos (n = 11) (Extended Data Fig. 1b,c). To identify the cell types that are represented in our dataset, we performed uniform manifold approximation and projection (UMAP) and Louvain clustering (Fig. 1d, Extended Data Fig.1d,e). We annotated the identified clusters based on a recent comprehensive scRNA-seq atlas of the developing mouse cerebral cortex^2^. We detected APs (expressing *Sox2, Pax6* and *Hes5*) as well as intermediate progenitors (IP; *Eomes, Neurog2, Btg2*) (Fig. 1e and Extended Data Fig. 1f). The progenitors formed a continuum in UMAP space with migrating neurons (MN; expressing *Nrp1*) and excitatory projection neurons (PN; *Neurod6, Tubb3*), including upper layer projection neurons (PN2; expressing *Satb2*). Although not derived from APs, we also captured a small proportion of interneurons (*Dlx1, Dlx2, Gad2*) and red blood cells (hemoglobin genes), but these cell types will not be the focus of this work and therefore excluded from further analysis. To exclude an IUE- or construct-dependent effect, we co-embedded our data with matched stages of the scRNA-seq atlas^2^ and compared cell type compositions. We observe strong overlap in UMAP space and comparable relative cell type abundances between our scDam&T-seq and the atlas data (Extended Data Fig. 1g-i).

We next inspected the LAD measurements and observed enrichment at known LAD regions across the genome (Fig. 1f and Extended Data Fig. 1j)^7^. Interestingly, aside from cell type-invariant constitutive LADs (cLADs), we also found both progenitor cell type-specific as well as neuron-specific LADs (dashed boxes; Fig. 1f). We next analyzed chromatin accessibility data as measured by Dam126 and observed a sharp enrichment at the promoters of expressed genes (Fig. 1g, Extended Data Fig. 1j-l). The Dam126 signal is primarily enriched on gene promoters according to their cell type-specific expression (Fig. 1g and Extended Data Fig. 1l). In addition, we compared the Dam126 scDam&T-seq data to scATAC-seq data, a commonly used assay for measuring chromatin accessibility^22^, of the same reference atlas that we used for validation of our scRNA-seq data^2^. The Dam126 data strongly correlates with the scATAC-seq data in a cell type-specific manner, both on individual genes and genome-wide (Extended Data Fig. 1l,m).

Together, we developed an efficient approach that combines IUE with scDam&T-seq, to enable joint measurement of transcriptomes and protein-DNA contacts in vivo in single cells of the developing mouse cerebral cortex.

### Widespread genome–lamina reorganization at long neuronal genes during neurogenesis

To dissect the interplay between spatiotemporal gene expression and genome–lamina contacts during neurogenesis, we inferred a pseudotime trajectory using Monocle 3^23^, excluding cells of non-cortical origin (Fig. 2a). We identified 677 genes that are dynamically expressed along the differentiation trajectory (Fig. 2b). Surprisingly, while LADs are generally gene-poor^7,13,24^, we found that more than half of these dynamically expressed genes (54%) contact the nuclear lamina in at least one of the cell types (henceforward referred to as dyn-LAD genes)(Fig. 2b). The remaining 46% of genes do not contact the nuclear lamina in any of the cell types and will therefore be referred to as dynamically expressed inter-LAD (dyn-iLAD) genes. Intrigued by the scale at which dynamically expressed genes contact the nuclear lamina during neurogenesis, we wondered about the biological functions of the dyn-LAD genes. Interestingly, while dyn-iLAD genes are enriched for biological processes related to cell proliferation and regulation of the cell cycle, dyn-LAD genes are highly enriched in functions that regulate neuronal processes (Fig. 2c and Extended Data Fig. 2a). These include the generation of neurons, neurogenesis and development of the nervous system more broadly (Fig. 2c and Extended Data Fig. 2a). We examined the nuclear lamina interactions for many such neuronal genes, and observed pronounced genome–lamina reorganization in both directions (i.e., away and towards the nuclear lamina) that can span megabase-sized domains encompassing such genes (Fig. 2d, Extended Data Fig. 2b). For example, *Satb2*, encoding a transcription factor involved in upper layer neuron specification and connectivity^28^, is activated during neurogenesis and detaches from the nuclear lamina (Fig. 2e). Conversely, the glutamate transporter transcript *Slc1a3*^29^, expressed specifically in APs, is deactivated and moves towards the nuclear lamina (Fig. 2e). We systematically examined how genome–lamina dynamics occur over the neurogenesis trajectory for all dyn-LAD genes. Almost all dyn-LAD genes showed a strong inverse relationship between their transcriptional activity and genome–lamina repositioning (Extended Data Fig. 2c,d). Genes that are deactivated over differentiation gradually move towards the nuclear lamina, while activating genes progressively move away (Extended Data Fig. 2c,d).

**Fig 2.**
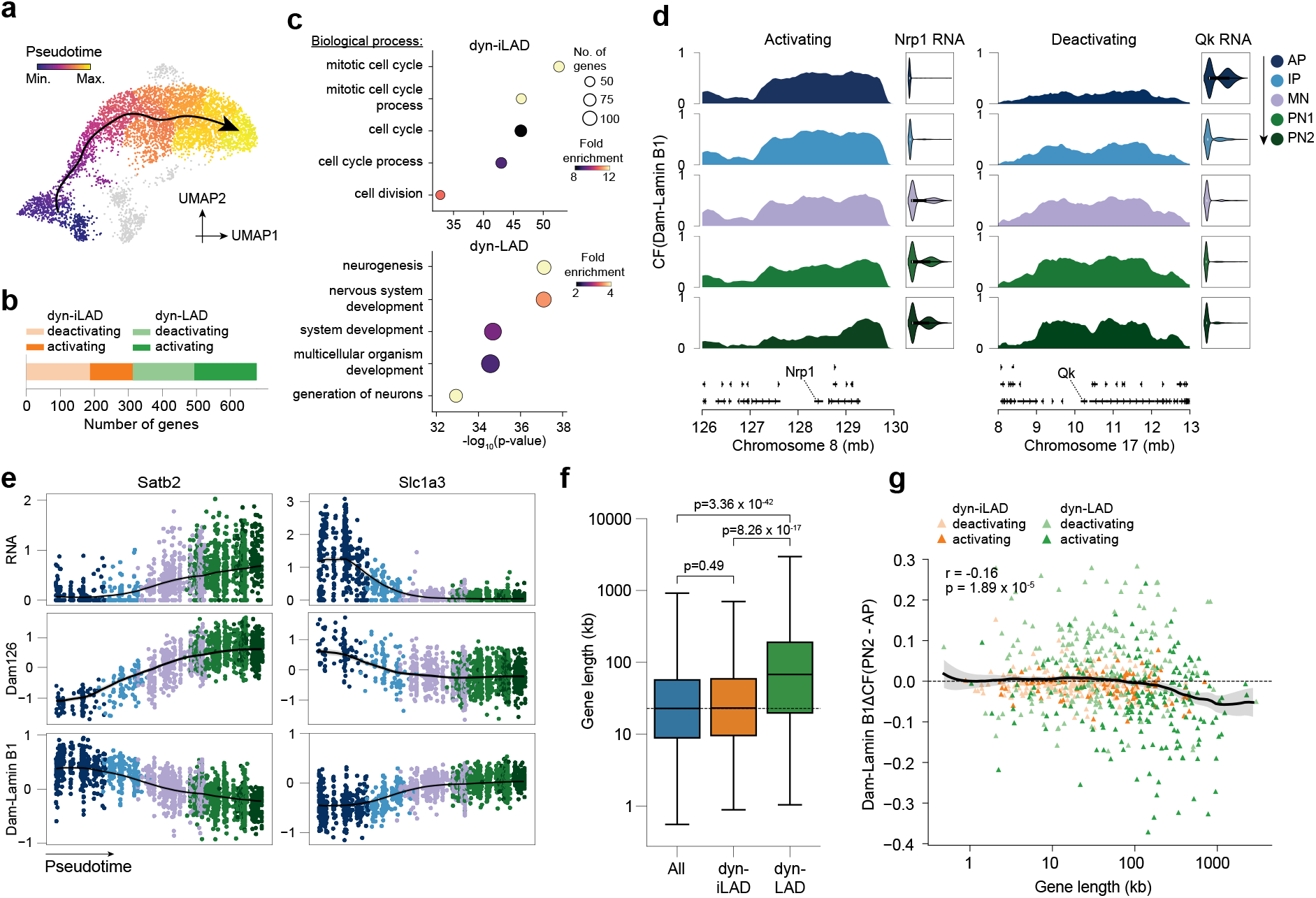
Prevalent genome–lamina reorganization at long neuronal genes during neurogenesis. **a**, UMAP plot with pseudotime^23^ projected on top. Arrow indicates direction of differentiation trajectory. **b**, Over half of the dynamically expressed (dyn) genes over pseudotime contact the nuclear lamina (LAD) in any of the neural cell types. **c**, Top 5 most significant terms of gene ontology analysis for biological processes on dyn-iLAD (top) and dyn-LAD (bottom) genes. **d**, LAD dynamics of representative dyn-LAD genes during neurogenesis. Cell type-specific Dam-Lamin B1 contract frequency (CF) is plotted in tracks and RNA expression in violins. **e**, Single-cell RNA expression, chromatin accessibility (Dam126) and nuclear lamina (Dam-Lamin B1) dynamics over pseudotime for representative dyn-LAD genes. **f**, Quantification of gene length for all expressed genes, dyn-iLAD and dyn-LAD genes. P-values from two-sided t-test. **g**, Nuclear lamina change of dyn-iLAD and dyn-LAD as a function of gene length. Pearson correlation (r) and p-value from a null hypothesis test.

To validate the observed inverse link between gene expression and genome–lamina proximity, we performed single-molecule RNA FISH (smFISH) for a selection of dyn-LAD genes on E16 cortical sections. We specifically designed probes targeting intronic regions, to simultaneously visualize nascent mRNA at the site of active transcription and the spatial position of the gene locus inside the nucleus^25,26^. As a control, we included probes targeting *Vim*, a gene that, according to our scDam&T-seq data, remains associated with the nuclear lamina in all cell types despite being highly expressed in APs—reflecting characteristics of previously described ‘escaper’ genes^15^. We consistently identified 1 to 2 nuclear foci per cell (Methods), which likely represent both the localization and activity of the two alleles (Extended Data Fig. 3a). We observed specific expression of the AP markers *Pax6* and *Hopx* of foci located in the nuclear interior of cells in the VZ, whereas the neuronal genes *Nav3, Nrp1* and *Satb2* were similarly detected, but in cells of the CP (Extended Data Fig. 3a). Moreover, the average spatial distance of genes to the nuclear periphery measured by smFISH strongly negatively correlates with their scDam&T-seq based nuclear lamina contact frequency (CF)(Extended Data Fig. 3b).

Interestingly, a subset of neuronal genes tends to be much longer than the average gene length and long genes have been implicated in neurodevelopmental and neurodegenerative disorders, including autism spectrum disorder (ASD) and Alzheimer’s disease^31–34^. We quantified the length of the above defined dyn-LAD genes and found that they are significantly longer than dyn-iLAD genes and the average gene length (Fig. 2f, Extended Data Fig. 4a). In addition, the longer the dyn-LAD gene, the more pronounced its genome–lamina repositioning during neurogenesis (Fig. 2g). As a result, we asked if gene length is more generally implicated in neural nuclear lamina association. We used Dam-Lamin B1 data of mouse embryonic stem cells (ESCs) as reference to identify neural specific genome–lamina interactions (Methods)^35^. We compared gene length for all protein-coding genes between LADs and inter-LADs (iLADs) of the different cell types and found that genes residing in neural LADs are significantly longer than genes in iLADs, or ESC LADs (Extended Data Fig. 4b). Moreover, the longer the gene; the higher the genome–lamina CF, suggesting a role for the nuclear lamina in regulating long genes in the developing cortex (Extended Data Fig. 4c).

Because long genes have been implicated in ASD^32^, which is thought to be a neurodevelopmental disorder, we cross-referenced the dyn-LAD genes with known ASD candidate genes described in the SFARI Gene database^79^. Notably, 20% (n = 74) of the dyn-LAD genes are known ASD candidate genes (Extended Data Fig. 4d), a proportion that is highly significant compared to chance (p = 2.4 × 10^-15^, hypergeometric test), as opposed to dyn-iLAD genes (n = 27, p = 0.3).

These results implicate prevalent genome–lamina reorganization in the regulation of long neuronal genes during cortical development.

### Neuronal genes are released from the nuclear lamina before transcriptional activation

Prompted by the scale of genome–lamina reorganization of neuronal genes during corticogenesis, we wondered if this spatial reorganization may be important to neuronal gene expression. To address this, we set out to dissect the sequence of gene regulatory events at the nuclear lamina during neurogenesis. Interestingly, a previous study in cell populations has suggested that some genes “unlock” from the nuclear lamina prior to transcriptional activation during in vitro differentiation of astrocytes^7^. However, generalizing the precise order of genome–lamina repositioning in relation to gene activity has remained largely elusive, in particular for in vivo developmental contexts.

Firstly, we examined both gene expression and genome–lamina dynamics along their shared pseudotime axis (Fig. 3a). We inspected genes that belong to the dyn-LAD class defined above (Fig. 2b), including deactivating genes expressed in APs, activating genes that are upregulated during neurogenesis and some rarer instances of transiently expressed genes (Fig. 3a). Importantly, to accurately determine the onset of transcriptional change, we quantified unprocessed pre-mRNA by distinguishing the proportion of unspliced from spliced mRNA reads. In our scDam&T-seq data, we detected 20% unspliced mRNA reads, similar to other protocols that rely on oligo-dT primers to capture polyadenylated mRNA molecules (Extended data Fig. 5a)^27^.

**Fig 3.**
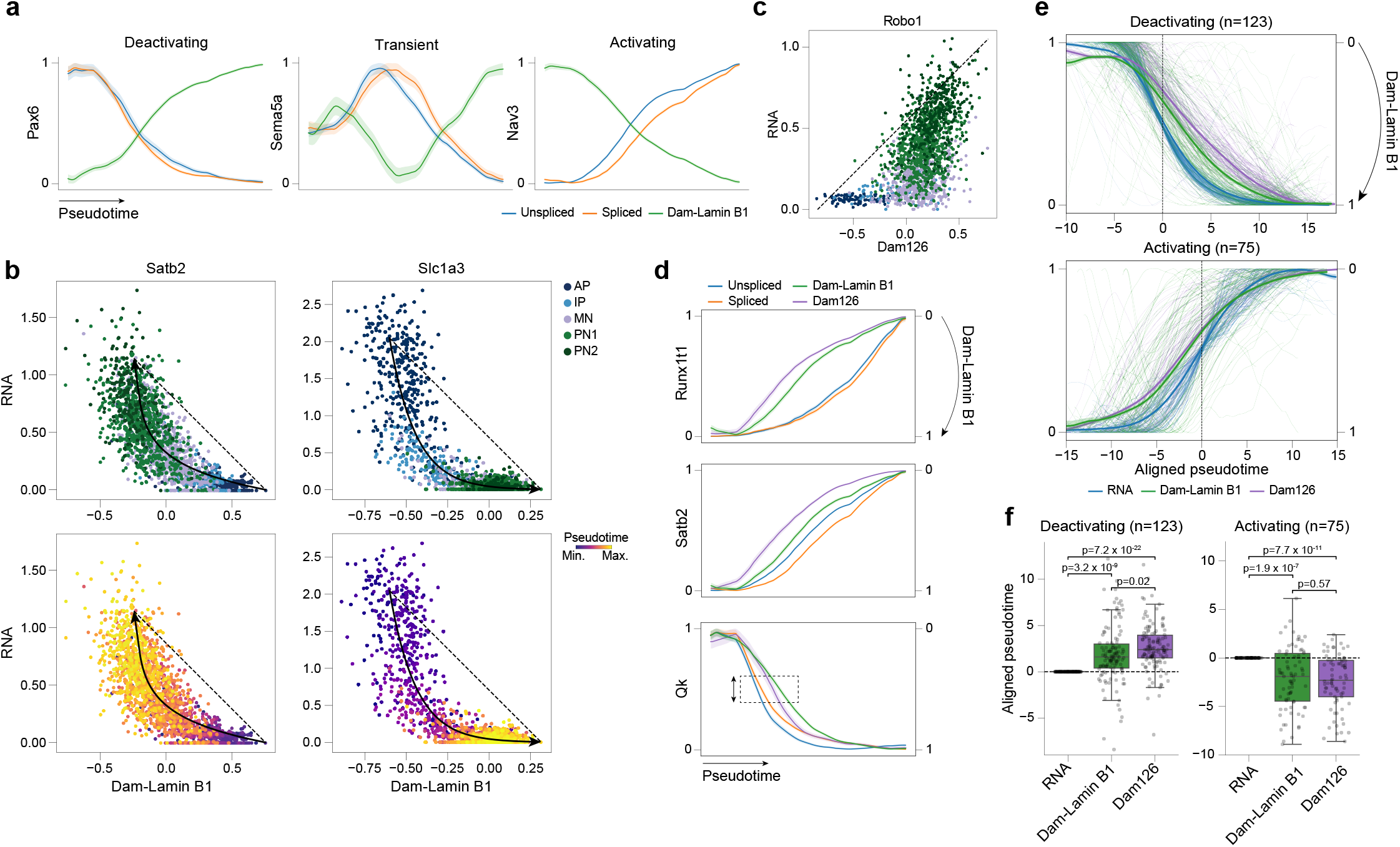
Long neuronal genes are released from the nuclear lamina before transcription activation. **a**, Dynamics of pre-mRNA (unspliced, blue), mature mRNA (spliced, orange) and genome–lamina contacts (Dam-Lamin B1, green) in a shared pseudotime space. Lines represent min-max scaled sliding window means with their 95% confidence interval in the shaded areas. **b**, Single-cell scatterplots of RNA expression and Dam-Lamin B1 for *Satb2* (left) and *Slc1a3* (right) colored by cell type (top) and pseudotime (bottom). The fitted lines result from a model that aims to explain the relationship between transcription and genome–lamina temporal dynamics (Methods). Arrowheads indicate direction of change. **c**, Single-cell scatterplot of RNA expression and chromatin accessibility (Dam126) dynamics of *Robo1* colored by cell type. Black interrupted line indicates the diagonal. **d**, Same as (**a**) with the addition of chromatin accessibility (Dam126, purple). Dam-Lamin B1 line is inverted for intuitive comparison of order of events. **e**, Clusters of similarly deactivating (top) or activating (bottom) genes (**Extended Data Fig. 3e**) plotted along aligned pseudotime (Methods). Genes are aligned based on expression (dashed line = 0). Each thin line represents a single gene and the thick lines the rolling mean of each modality with its 95% confidence interval. **f**, Quantification of aligned pseudotime (Methods) moments where it reaches the middle 20% of its dynamic range (i.e., between the 40th and 60th percentile, box in (**d**)) of the dynamics range for clusters of similarly activating and deactivating genes. P-values from two-sided t-test.

Although genome–lamina and gene expression dynamics are roughly comparable along pseudotime, our multimodal data offers a higher resolution of temporal dynamics, particularly relevant when events are closely related in time (Fig. 3a). To this end, we directly plotted both readouts of our scDam&T-seq data for representative dyn-LAD genes (Fig. 3b, Extended Data Fig. 5b). Interestingly, the cells in these plots showed a highly non-random distribution, revealing a temporal relationship between gene expression and genome–lamina dynamics. Alignment along the diagonal would indicate that both events are temporally coordinated, but we observe clear divergence from the diagonal, suggesting sequentiality. For example, *Satb2*, a neuronal gene that is 184.7 kb long, first spatially repositions away from the nuclear lamina before upregulation of expression (Fig. 3b). In contrast, *Slc1a3* (76.5 kb), is deactivated as cells differentiate and transcription downregulation happens before the locus moves towards the nuclear lamina (Fig. 3b). We inspected a range of dyn-LAD genes in this manner and observed a more general principle of activating genes that first detach from the nuclear lamina before upregulation and deactivating genes that are downregulated before moving towards the nuclear lamina (Extended Data Fig. 5b,c).

It has been proposed that changes in chromatin accessibility may prime cells for lineage commitment by preceding gene expression^30^. We wondered how chromatin accessibility dynamics relate to genome–lamina reorganization during neurogenesis. To address this, we further examined the Dam126 scDam&T-seq data. In agreement with previous findings of shared scRNA and scATAC-seq on mouse skin tissue^30^, chromatin accessibility measured by Dam126 mainly precedes gene expression during neurogenesis (Fig. 3c, Extended Data Fig. 5d). For genes that are deactivated, chromatin compaction of the locus rather happens after downregulation of expression (Extended Data Fig. 5d). To obtain an integrative view of the order of gene regulatory events at the nuclear lamina, we quantified the onset of chromatin accessibility, gene expression and genome– lamina change along pseudotime for selected hierarchical clusters of similarly activating or deactivating dyn-LAD genes, such as *Runx1t1* (150.2 kb, activating), *Satb2* (184.7 kb, activating) and *Qk* (112.8 kb, deactivating)(Fig. 3d,e and Extended Data Fig. 5e). Of note, we inverted the Dam-Lamin B1 axis for visualization purposes to enable intuitive comparison of temporal dynamics (Fig. 3d,e). For deactivating dyn-LAD genes, we generally observed downregulation of transcription as the first event, before relocation of the locus towards the nuclear lamina and subsequent chromatin compaction (Fig. 3e,f). For activating dyn-LAD genes, we found instances of long neuronal genes where chromatin first decompacts before the locus is released from the lamina after which transcription activation takes place (Fig. 3d). Nevertheless, on average, we found that for activating dyn-LAD genes: chromatin becoming accessible and genome–lamina repositioning happening almost coincidentally, before expression of the gene is upregulated (Fig. 3e,f).

Collectively, these results identify genome–lamina repositioning as an early event along the regulatory cascade of long neuronal genes, suggesting that genome–lamina repositioning is not simply a consequence of, but may be required to enable neuronal gene expression.

### MeCP2 binds long nuclear lamina regulated genes before release and transcriptional activation

Intrigued by our finding that genome–lamina reorganization during corticogenesis is strongly biased towards long genes, we next wondered what factors may be involved in this process. Interestingly, the X-chromosome encoded methyl CpG binding protein 2 (MeCP2) has previously been found to repress the expression of long genes in the brain^31,38^. Mutations in MeCP2 cause Rett syndrome (RTT), a severe neurodevelopmental disorder with features of ASD, mainly affecting females^16,43^. MeCP2 is highly upregulated in maturing neurons^42^, but is also expressed in the developing cortex and upregulated during neurogenesis (Extended Data Fig. 6a), albeit at lower levels^43–45^. Although overt RTT-related symptoms typically emerge postnatally, the severe phenotypes observed in *Mecp2*-null male mice and the rarity of human male RTT births suggest that MeCP2 function is already important during embryonic development^43,48,49^. Indeed, several studies have uncovered transcriptional and cellular defects resulting from MeCP2 disruption during neurogenesis in both mouse and human^44,45,50,51^. Because our results uncovered that genome– lamina reorganization during neurogenesis involves long genes, we hypothesized whether there is a link between nuclear lamina and MeCP2 regulation of such genes. To this end, we set out to map MeCP2 binding profiles in single cells of the developing mouse cortex with scDam&T-seq.

Firstly, we designed a DamID construct that fuses MeCP2 to Dam126 to perform IUE combined with scDam&T-seq on E14 stage embryos (Fig. 4a). Using immunofluorescence imaging, we examined the expression of the MeCP2-Dam126 fusion protein across cortical layers and validated its localization inside the nucleus (Fig. 4b). MeCP2-Dam126 specifically localized to DNA dense regions, consistent with previous studies that visualized MeCP2 localization^52,53^. Next, we dissociated cortical tissue to perform scDam&T-seq on FACS sorted mRuby+ cells. We obtained high quality single-cell joint MeCP2 binding and transcriptomic data (Extended Data Fig. 6b,c). To verify the localization of MeCP2-Dam126 signal along the genome, we correlated the data to MeCP2 ChIP-seq data of the maturing cortex and observed that they strongly agree (Extended Data Fig. 6c,d)^46^. Moreover, MeCP2-Dam126 signal is specifically enriched on genes that are misregulated in the maturing cortex of MeCP2-deficient mice (Extended Data Fig. 6e)^46^. We verified the integration of the MeCP2-Dam126 scDam&T-seq data with the Dam-Lamin B1 and Dam126 cells by projecting the MeCP2-Dam126 cells on the above defined UMAP (Fig. 1d), which was used as a reference (Fig. 4c, Extended Data Fig. 6f). Together, these results establish IUE combined with scDam&T-seq as a robust approach to dissect the gene regulatory function of MeCP2 on transcription at single-cell resolution during cortical development.

**Fig 4.**
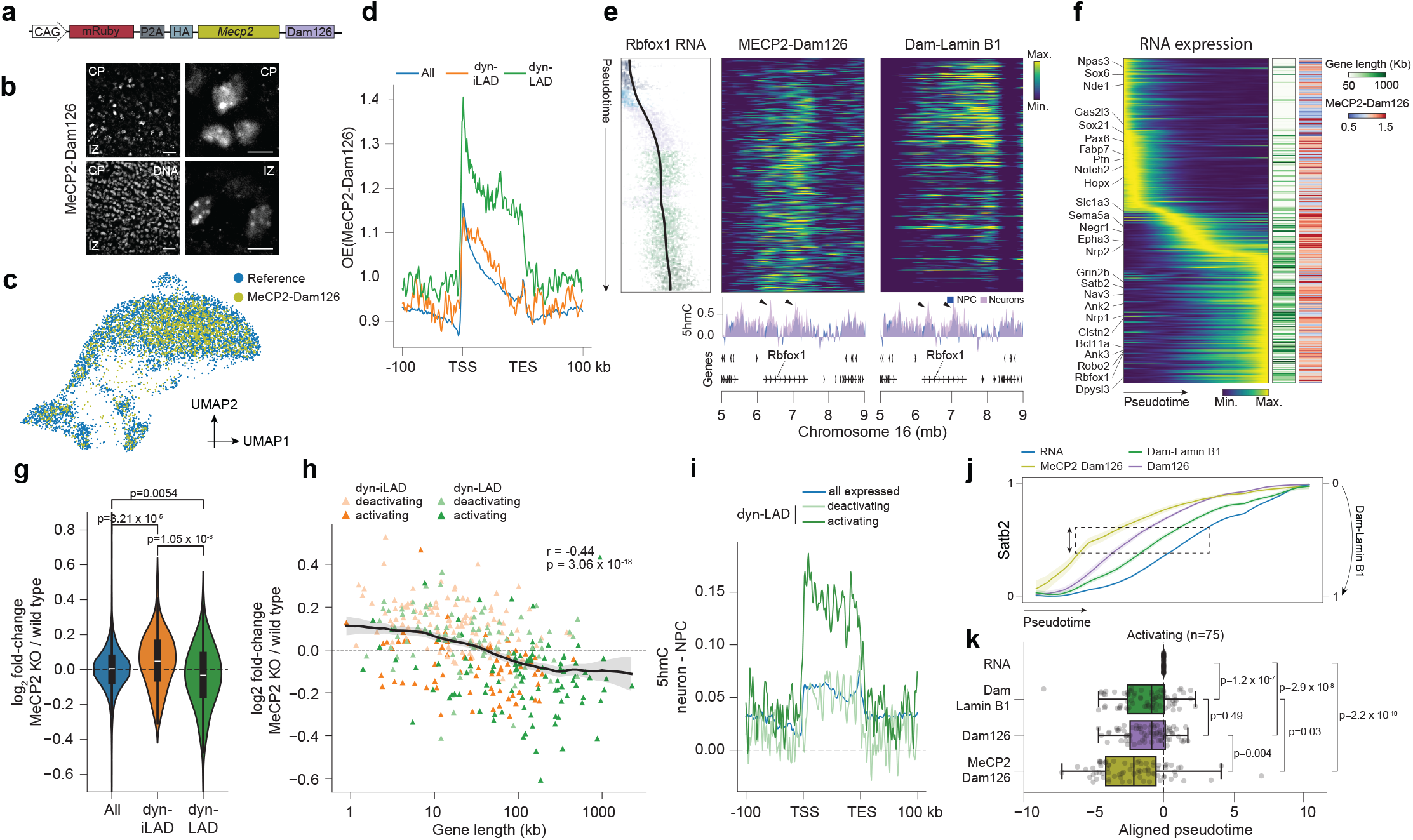
MeCP2 binds long nuclear lamina regulated genes before release and transcriptional activation. **a**, MeCP2 DamID construct design. **b**, Immunofluorescence on HA-tag of the MeCP2-Dam126 construct on cortical tissue slides at E16. Bottom left image shows a DAPI DNA staining. Left images are 20X magnifications and white scale bar is 25 µm, and right are 60X magnifications and white scale bar is 5 µm. **c**, MeCP2-Dam126 cells projected on a reference UMAP plot. **d**, MeCP2-Dam126 signal aligned over all expressed genes, dyn-iLAD and dyn-LAD genes. **e**, Dynamics of *Rbfox1* over neurogenesis. Below 5hmC enrichment in neural progenitor cells (NPC, blue) and neurons (purple) of E15.5 mouse embryos as measured by hMeDIP^36^. **f**, RNA expression heatmap of dyn-LAD genes ordered on their peak of expression. Colorbars depict gene length (left), MeCP2-Dam126 (right). **g**,**h**, Gene expression changes measured by microarray of the cortex from MeCP2 KO (n = 3) and WT control (n = 3) E15.5 mouse embryos^44^ for all expressed genes, dyn-iLAD and dyn-LAD genes in violin plot (**g**) or as function of gene length (**h**). P-values from two-sided t-test. **i**, The difference of 5hmC levels between NPCs and neurons of E15.5 mouse embryos measured by hMeDIP^36^ on activating and deactivating dyn-LAD genes. **j**, *Satb2* dynamics. Lines represent scaled sliding window means with 95% confidence interval in the shaded area. Dam-Lamin B1 line is inverted. **k**, Quantification of aligned pseudotime (box in (**j**); Methods) of similarly activating genes (**Extended Data Fig. 5e**).

Interestingly, we found that MeCP2 is highly bound to the bodies of dyn-LAD genes during neurogenesis (Fig. 4d), which is not explained by chromatin accessibility (Extended Data Fig. 7a-c). In addition, we observed that MeCP2 binds dyn-LAD genes in a length-dependent manner, where the longest genes display the highest level of MeCP2 binding, relative to shorter genes (Extended Data Fig. 7d,e). Because MeCP2 regulation of long genes has mostly been studied the maturing cortex of postnatal mice^31,38,46^, we re-analyzed MeCP2 ChIP-seq data^46^ of that stage and found that there too MeCP2 is specifically bound to dyn-LAD genes (Extended Data Fig. 7f). This suggests that MeCP2 targets the same nuclear lamina regulated long genes during pre- and postnatal cortical development. However, in the maturing cortex of postnatal mice, MeCP2 represses the expression of long genes^31,38^, while our results uncovered that MeCP2 binds these same genes during neurogenesis, a period during which they are robustly activated and detach from the nuclear lamina (Fig. 4e,f). Of note, although MeCP2 is primarily described as a transcriptional repressor, studies have also reported its role in transcriptional activation^39–41^. To investigate this apparent discrepancy, we analyzed transcriptomic data of both the maturing (postnatal week 8)^46^ and embryonic (E15.5)^44^ cortex of MeCP2 knockout (KO) mice in comparison to wildtype (WT) control mice. Interestingly, while dyn-LAD genes are upregulated in the maturing cortex of MeCP2 KO mice (Extended Data Fig. 7g-i), they are downregulated in the embryonic cortex of MeCP2 KO mice in a gene length-dependent manner (Fig. 4g,h). Furthermore, as opposed to being globally upregulated in the maturing cortex of MeCP2 KO mice^31,38^, we found that long genes are globally downregulated in the cortex of MeCP2 KO mouse embryos (Extended Data Fig. 7j). These findings suggest that while MeCP2 represses long nuclear lamina regulated genes in the postnatal cortex, it activates them in the prenatal cortex.

We wondered what molecular mechanisms may underlie this apparent developmental context-specific effect of MeCP2 on dyn-LAD genes. MeCP2 has strong affinity for methylated DNA^31,37,40^ and DNA methylation is highly dynamic during mammalian brain development^47^. It has previously been shown that MeCP2 represses long gene expression by binding to methylated CA (5mCA) sites within such genes^31^, and levels of 5mCA, deposited by the methyltransferase DNMT3A, increase in the brain after birth^47^. We therefore analyzed bisulfate sequencing data of the maturing cortex^46^ and observed that 5mCA is specifically enriched on dyn-LAD genes at that stage (Extended Data Fig. 8a,b). In contrast, an epigenetic modification that accumulates during neurogenesis in the developing brain is 5-hydroxymethylcytosine (5hmC)^36^, resulting from oxidation of 5-methylcytosine (5mC) by TET enzymes^54,55^. Interestingly, MeCP2 has strong affinity for 5hmC and this mark is highly associated with active genes in the brain^40^. Consequently, we set out to investigate levels of 5hmC on nuclear lamina regulated genes during neurogenesis^36^. Along with gradual upregulation of TET protein gene expression (Extended Data Fig. 8c), we observed that 5hmC specifically accumulates on dyn-LAD genes that are activated during neurogenesis (Fig. 4e,i). In addition, 5hmC accumulates on dyn-LAD genes in a length-dependent manner, where the longer genes accumulate the strongest levels of 5hmC during neurogenesis (Extended Data Fig. 8d), with a concomitant loss of 5mCG (Extended Data Fig. 8e)^80^. These findings suggest that the developmental context-specific effect of MeCP2 that we observed on long nuclear lamina regulated genes may be mediated by the methylation status of such genes.

Finally, as we observed earlier that dyn-LAD genes detach from the nuclear lamina prior to their transcriptional activation, we next asked when MeCP2 acts within the regulatory cascade that leads to the activation of such long lamina regulated genes. We found that MeCP2 binds dyn-LAD genes prior to their release from the nuclear lamina and before these gene loci become accessible, followed by their transcriptional activation (Fig. 4i,j).

In sum, our findings suggest a link between the nuclear lamina and MeCP2 in the regulation of long genes during neurogenesis. MeCP2 may be involved in preparing hydroxymethylated long genes for genome–lamina detachment and subsequent transcriptional activation. In addition, our analyses suggest that MeCP2 may act as a context-specific regulator, capable of both repressing and activating long genes depending on developmental stage and associated methylation status of these genes—warranting further studies to elucidate the mechanisms underlying this dual functionality.

## Discussion

In this study, we developed an approach that combines in utero electroporation with single-cell DamID and transcriptome sequencing (scDam&T-seq) to profile genome–lamina interactions in vivo in the developing mouse cerebral cortex.

Although we mainly focused on LADs, our approach is broadly applicable to any protein of interest fused to Dam. Indeed, many Dam fusion proteins have already been developed and validated for profiling histone post-translational histone modifications, transcription factors and DNA repair proteins^56–59^. Furthermore, by employing a catalytically impaired Dam126 mutant^20^, we achieved highly specific single-cell chromatin accessibility profiles in the developing cortex.

Using this approach, we found that hundreds of neuronal genes undergo genome–lamina repositioning during neurogenesis. This observation aligns with recent bulk data in humans showing widespread spatial reorganization of neuronal genes during cortical development, suggesting a conserved mechanism across mammals^60^. Importantly, our results uncovered that nuclear lamina regulated neuronal genes during neurogenesis are notably long (≥100 kb). Long genes have been implicated in diverse physiological and neurological contexts, including aging^33^, ASD^32^ and Alzheimer’s disease^33^—and our results reveal a previously unrecognized regulatory step in their regulation in the developing brain, marked by extensive genome–lamina reorganization.

A longstanding question in LAD biology is whether genome–lamina repositioning is a cause or consequence of transcription^61^. Previous studies addressed this using engineered locus-repositioning assays in cell lines^12,15^. By leveraging our single-cell data, we found that neuronal genes are released from the lamina prior to transcriptional activation during in vivo development of the mammalian cortex^7^. Conversely, genes expressed in neural stem cells become transcriptionally repressed before repositioning towards the nuclear lamina. These findings suggest that while nuclear lamina association may not initiate gene repression, detachment from the nuclear lamina appears to be a necessary precondition for neuronal gene activation.

Our analyses suggest a link between genome-lamina reorganization at long neuronal genes and the transcriptional regulator methyl binding protein MeCP2^31,38^ during neurogenesis. We showed that MeCP2 specifically accumulates on long nuclear lamina regulated genes and loss of MeCP2 disrupts their expression in the developing cortex. These findings align with reports of early-onset deficits that precede overt Rett syndrome symptoms^44,45,50,51^. Notably, we found that while long nuclear lamina regulated genes are repressed by MeCP2 in the maturing cortex^31,38^, MeCP2 regulates these same genes during neurogenesis, when these genes are released from the nuclear lamina and robustly activated. These findings address a longstanding paradox regarding MeCP2’s dual role in transcriptional repression and activation^39–41^, by suggesting that it has stage-specific functions during development. We showed that a potential molecular mechanism through which MeCP2 exerts this developmental stage-specific effect on long nuclear lamina regulated genes, is the DNA methylation state of such genes. Notably, 5hmC emerges as a compelling candidate, because it accumulates on nuclear lamina regulated genes during neurogenesis in a length-dependent manner—and MeCP2 has strong affinity for this epigenetic modification^20^. In addition, our finding that MeCP2 activates long nuclear lamina regulated genes during neurogenesis aligns with the strong association of 5hmC with active genes in the brain^40^.

Interestingly, our single-cell MeCP2 data suggests that MeCP2 binds to hydroxymethylated long genes before their release from the nuclear lamina, supporting a model in which MeCP2 plays a role in the genome-lamina detachment of such genes and their subsequent transcriptional activation during neurogenesis (Fig. 5). In support of this, a previous study found that MeCP2 is responsible for nucleosome decondensation of hydroxymethylated CA repeats inside LADs in mouse fibroblasts^65^. In addition, a recently published report employing DNA-RNA MERFISH on the mouse cortex identified a radial-position-dependent DNA reorganization in excitatory neurons of MeCP2-deficient mice^66^. It has previously been shown that MeCP2 associates with the transcriptional activator CREB1 to activate gene expression^43^, and its function is critical for neurogenesis^67^. We speculate that during neurogenesis, MeCP2 plays a role in priming long genes for nuclear lamina release by associating with CREB1 that in turn results in the acetylation of histone tails^68^, a post-translational modification that has been demonstrated to antagonize LADs^9^.

**Fig 5.**
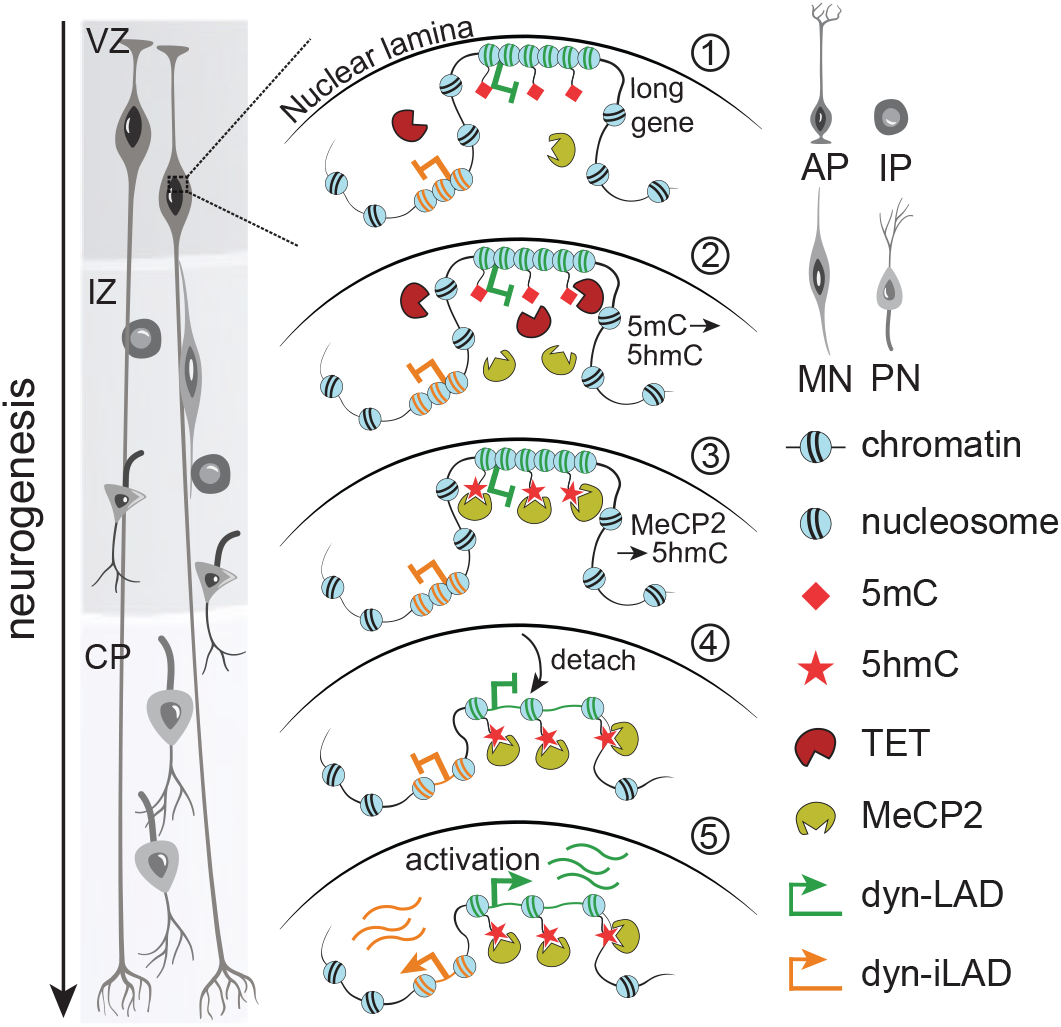
Model depicting the cascade of long gene activation during neurogenesis. 1, Long genes reside in neural specific 5mC marked LADs. 2, During neurogenesis, expression of TET enzymes and MeCP2 is upregulated. 3, TET enzymes oxidise 5mC to 5hmC on dyn-LAD genes that will be activated and MeCP2 binds. 4, Activating dyn-LAD genes chromatin becomes accessible and they are released from the lamina. 5, At the end of the gene regulatory cascade, transcription of the genes is activated. Cartoon partially adapted from ref. 78.

Taken together, we provide an experimental framework to study gene regulatory mechanisms in single cells of the developing mouse brain. Using this framework, we identified prevalent genome–lamina reorganization that precedes long neuronal gene activation during neurogenesis, that is potentially mediated by MeCP2. These findings point towards the intriguing possibility that MeCP2 can act as a context-dependent regulator of long genes by influencing genome–lamina interactions, offering new insights into its role in neurodevelopment and the etiology of Rett syndrome.

**Extended Data Figure 1.**
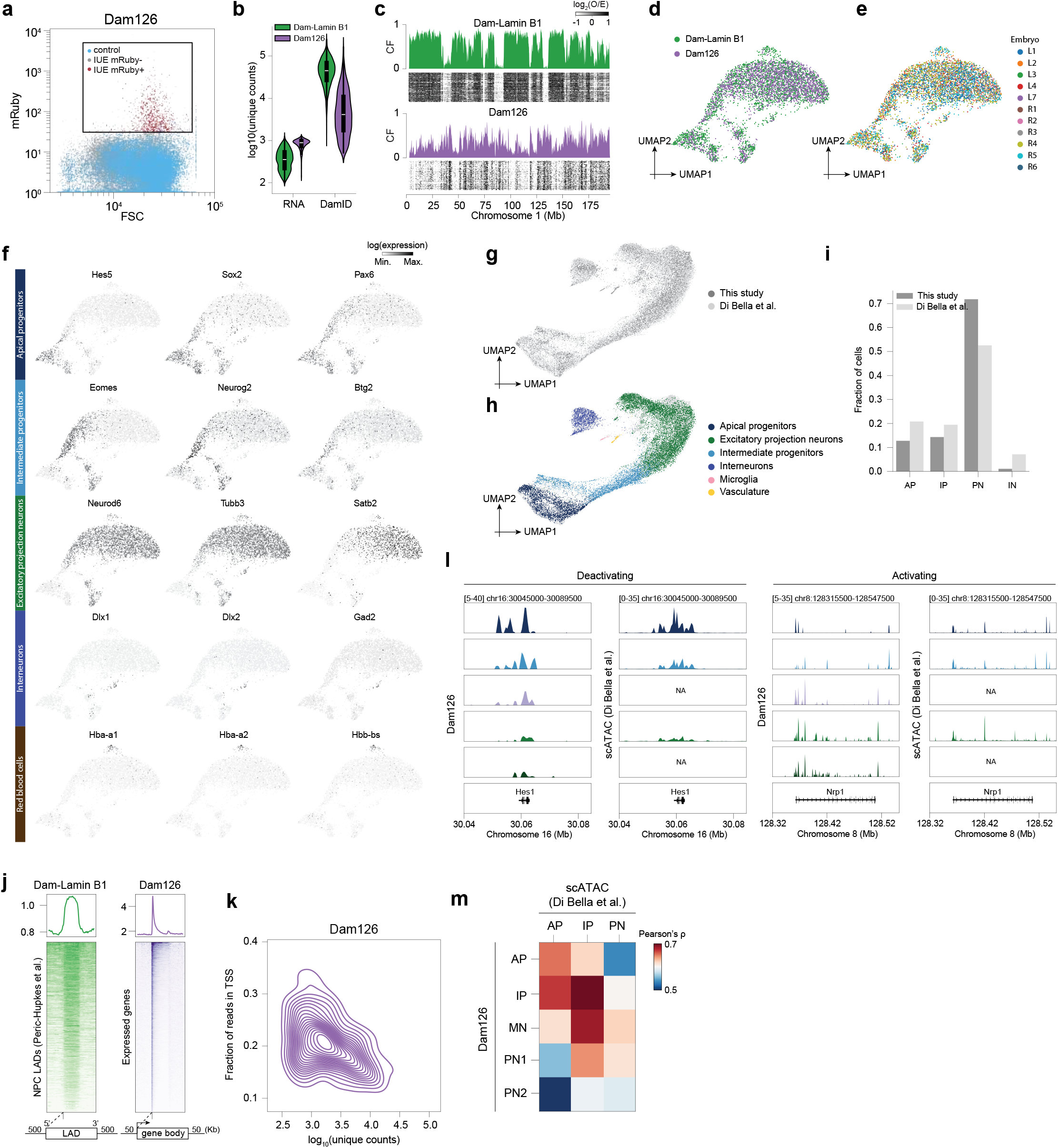
Validation of in utero electroporation scDam&T-seq transcriptome, LAD and chromatin accessibility data in the developing mouse cortex. **a**, FACS analysis on dissociated cortical tissue after IUE with Dam126. mRuby+ cells were sorted for scDam&T-seq (Methods). **b**, Unique transcripts and DamID reads for both constructs. **c**, Single-cell heatmaps for chromosome 1 for both constructs. **d**,**e**, UMAP plots with distribution of constructs (**d**) and embryos (**e**). **f**, UMAP plots with cell type-specific marker expression. **g**,**h**, UMAP co-embedding of scRNA-seq of atlas dataset^2^ and scDam&T-seq data (**g**) and with annotated cell types of the atlas dataset (**h**). **i**, Comparison of cell type fractions between atlas^2^ dataset and scDam&T-seq data. **j**, Dam-Lamin B1 data aligned over neural progenitor cell (NPC) LADs (left)^7^ and Dam126 over expressed genes (right). **k**, Distribution of unique Dam126 reads against fraction of reads overlapping TSS (2 kb flanking window). **l**, Cell type-specific chromatin accessibility on marker gene loci as measured by Dam126 (left) and scATAC-seq (right)^2^. **m**, Cell type-specific genome-wide Pearson correlation of chromatin accessibility measured by Dam126 and scATAC-seq^2^. 100 kb consecutive genomic bins were used.

**Extended Data Figure 2.**
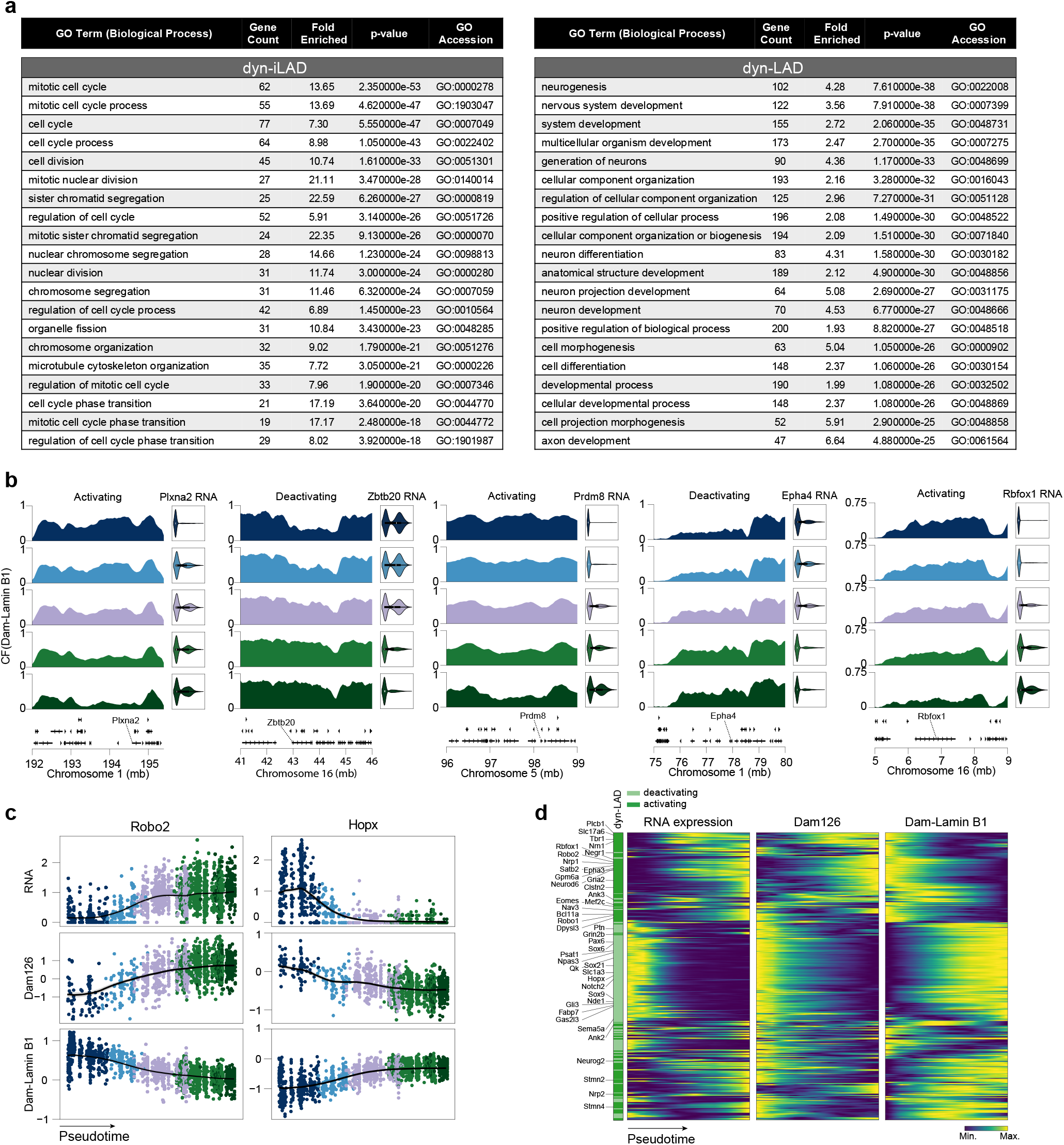
Widespread genome-lamina reorganization at neuronal genes. **a**, Top 20 most significant terms of gene ontology analysis for biological processes on dyn-iLAD (left) and dyn-LAD (right) genes. **b**, Five representative dyn-LAD genes (**Fig. 2b**). Cell type-specific Dam-Lamin B1 contract frequency (CF) is plotted in tracks and RNA expression in violins. **c**, Single-cell RNA expression, chromatin accessibility (Dam126), nuclear lamina (Dam-Lamin B1) dynamics over pseudotime of representative dyn-LAD genes. Black lines represent the sliding window means with 95% confidence interval in the shaded area. **d**, Heatmaps showing hierarchical clustering of RNA expression (left), chromatin accessibility (Dam126, middle) and genome–lamina (Dam-Lamin B1, right) dynamics over pseudotime of all dyn-LAD genes.

**Extended Data Figure 3.**
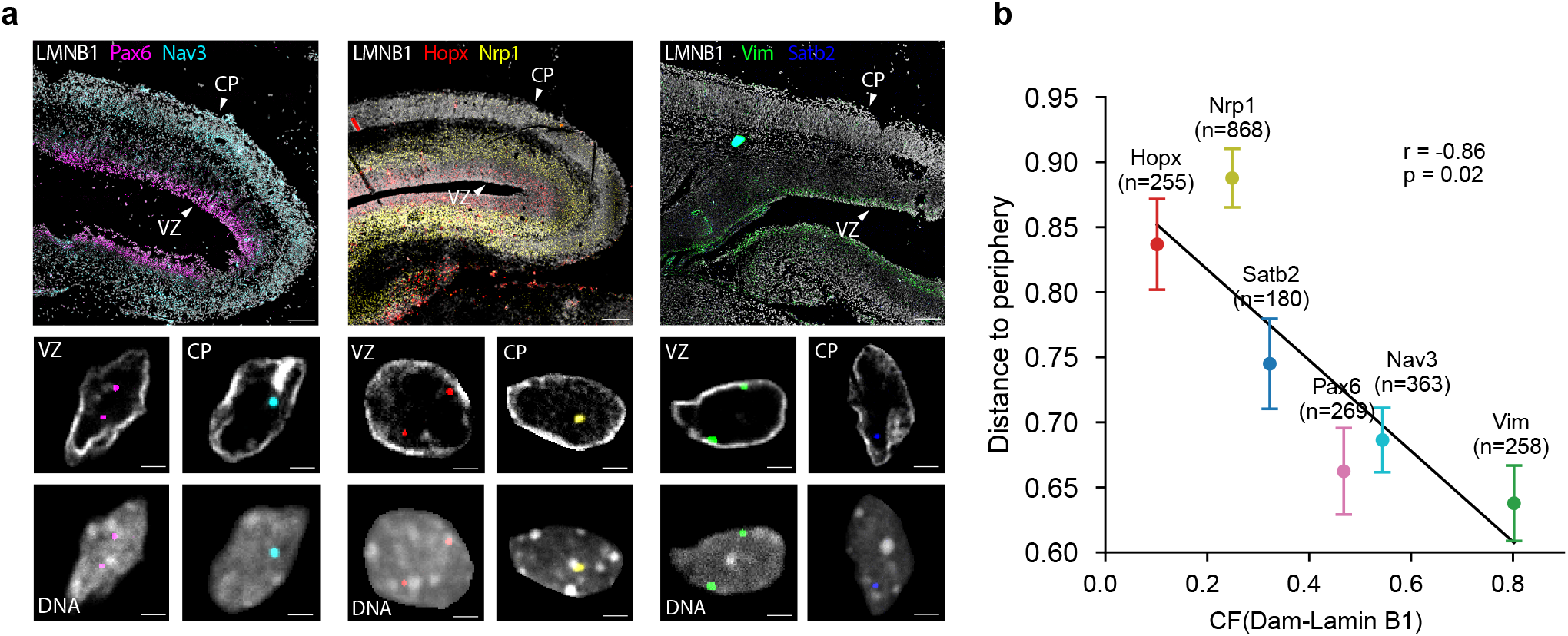
Spatial intron FISH identifies radial distance of transcribed nuclear lamina regulated genes. **a**, Intron-specific RNA FISH images of E16 cortical tissue slides for probes *Pax6* (magenta), *Nav3* (cyan), *Hopx* (red), *Nrp1* (yellow), *Vim* (green) and *Satb2* (blue). Top images are 20X magnifications, white scale bar is 50 µm. Bottom images are 100X zooms of the top image, white scale bar is 2 µm. **b**, Correlation between distance to periphery quantified from intron RNA FISH (Methods) and Dam-Lamin B1 contact frequency (CF). Black line represents a linear regression fit. Pearson correlation (r) and p-value from a null hypothesis test are shown.

**Extended Data Figure 4.**
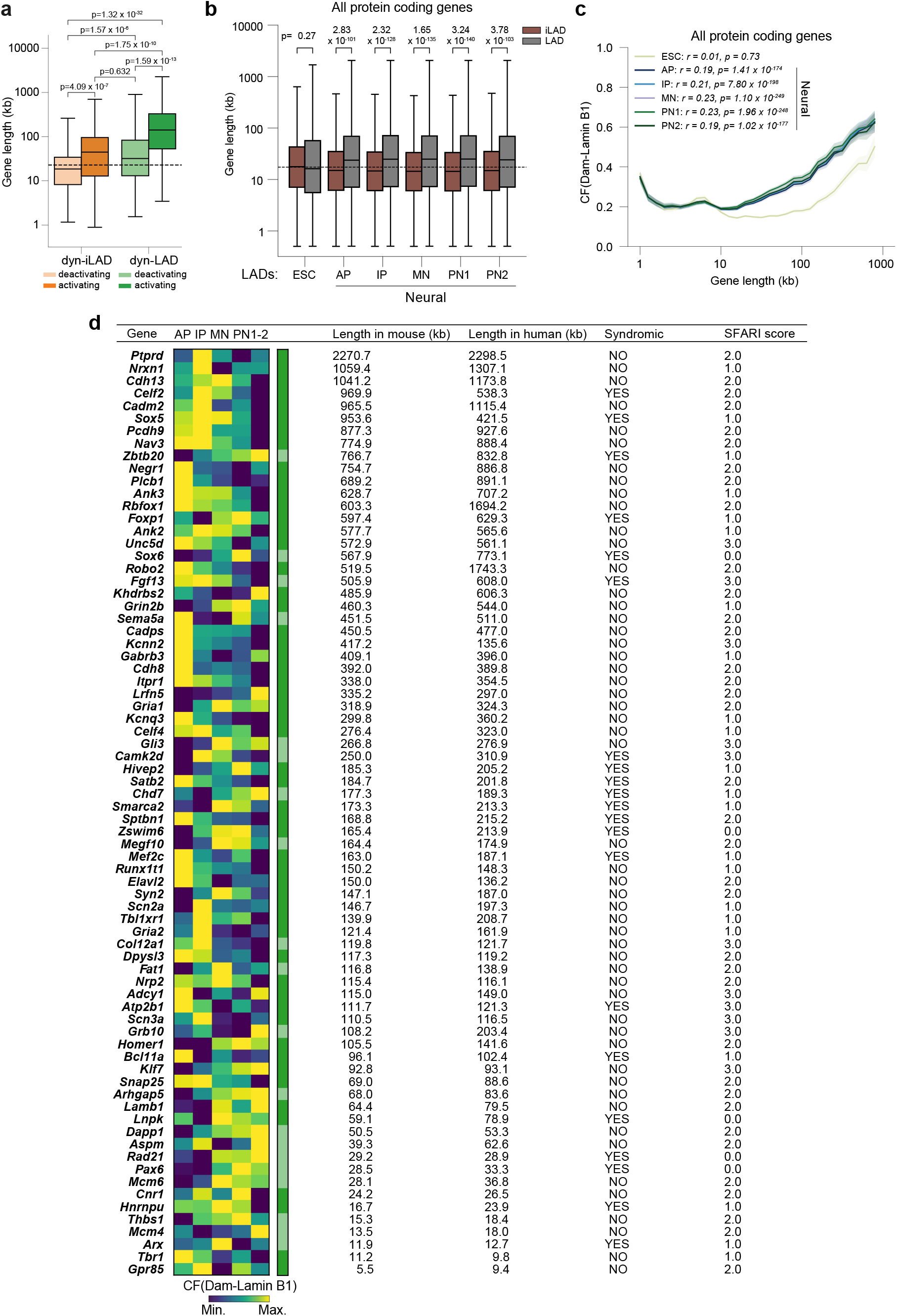
Long nuclear lamina regulated genes are associated with ASD risk. **a**, Quantification of gene length for dyn-iLAD and dyn-LAD genes stratified by deactivation or activation during neurogenesis. The dashed line is the median length of all expressed genes. P-values from two-sided t-test. **b**, Gene length for all protein coding genes stratified by whether their promoter (i.e. TSS bin) resides in LADs or not (iLAD) in the defined cell types. P-values from two-sided t-test. **c**, Dam-Lamin B1 contact frequency (CF) as a function of gene length for all protein coding gene promoters (i.e. TSS bin) split over the different cell types. Lines represent sliding window means with 95% confidence interval in the shaded area. **d**, 74 dyn-LAD genes overlapping with ASD candidate genes (*P* = 2.4 × 10^-15^, hypergeometric test) of the SFARI Gene database^79^, ordered by their length. Heatmap is scaled Dam-Lamin B1 CF for neural cell types.

**Extended Data Figure 5.**
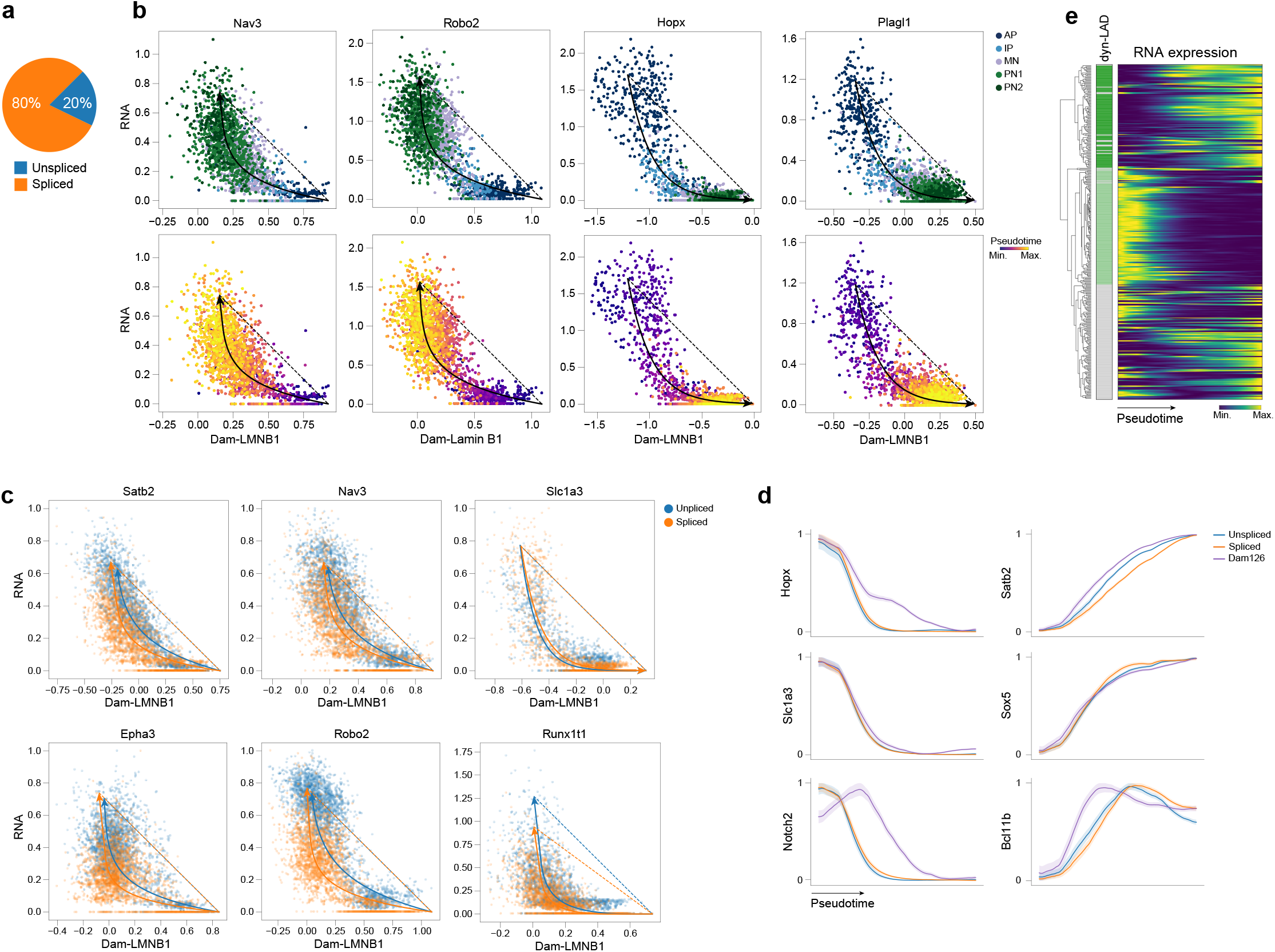
The sequence of gene regulatory events at the nuclear lamina during neurogenesis. **a**, Pie chart with fraction of spliced and unspliced mRNA reads detected by in utero electroporation scDam&T-seq. **b**, Single-cell scatterplots of RNA expression and Dam-Lamin B1 dynamics for four representative genes colored by cell type (top) and pseudotime (bottom). The fitted lines result from a model that aims to explain the relationship between transcription and genome–lamina temporal dynamics (Methods). Arrowheads indicate direction of change. **c**, Same as (**b**) split over pre-mRNA (unspliced, blue) and mature mRNA (spliced, orange) levels for 6 representative genes. **d**, Dynamics of pre-mRNA (unspliced, blue), mature mRNA (spliced, orange) and chromatin accessibility (Dam126, purple) in a shared pseudotime space. Lines represent min-max scaled sliding window means with their 95% confidence interval in the shaded areas. **e**, Heatmap showing hierarchical clustering of RNA expression dynamics over pseudotime of all differentially expressed LAD genes. Selected hierarchical clusters of similarly activating (dark green) or deactivating (light green) genes are depicted in the color bar.

**Extended Data Figure 6.**
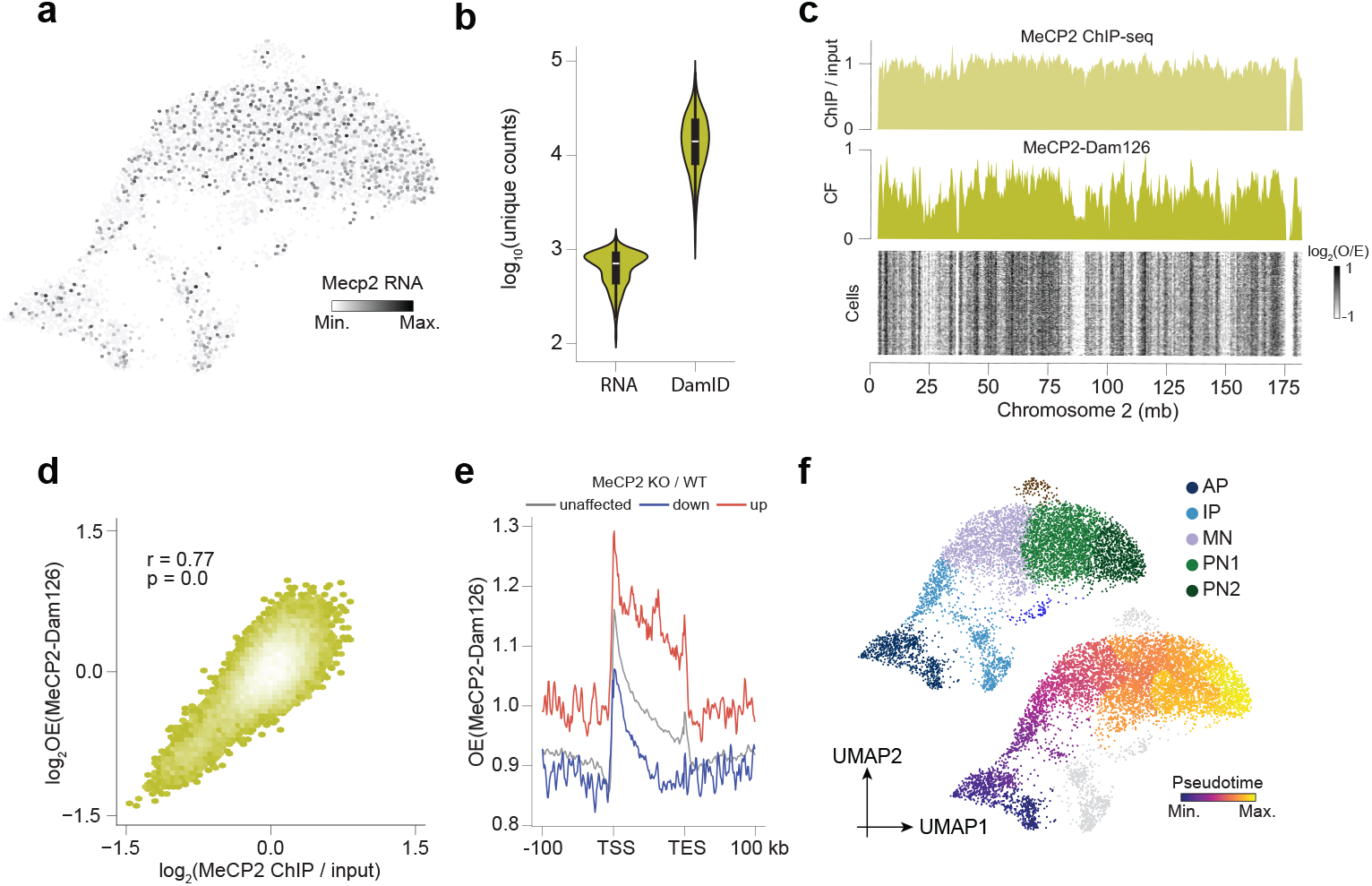
scDam&T-seq jointly maps MeCP2 binding and transcription in single cells of the developing mouse cortex. **a**, UMAP plot colored by *Mecp2* gene expression. **b**, Unique transcripts and MeCP2-Dam126 reads in single cells. **c**, Single-cell heatmap along chromosome 2 for MeCP2-Dam126 cells with contact frequency (CF) above. Top track is of publicly available MeCP2 ChIP-seq of the mouse cortex^46^. **d**, Genome-wide correlation between MeCP2-Dam126 and MeCP2 ChIP-seq^46^ signal in 100 kb genomic bins. Pearson correlation (r) and p-value from null hypothesis test are shown. **e**, MeCP2-Dam126 signal aligned over MeCP2 target genes defined in RNA-seq data from the maturing cortex of MeCP2 KO mice^46^. **f**, MeCP2-Dam126 cells projected on a reference UMAP plot as defined in **Fig. 1d**, colored by cell type (top) and pseudotime (bottom)^23^.

**Extended Data Figure 7.**
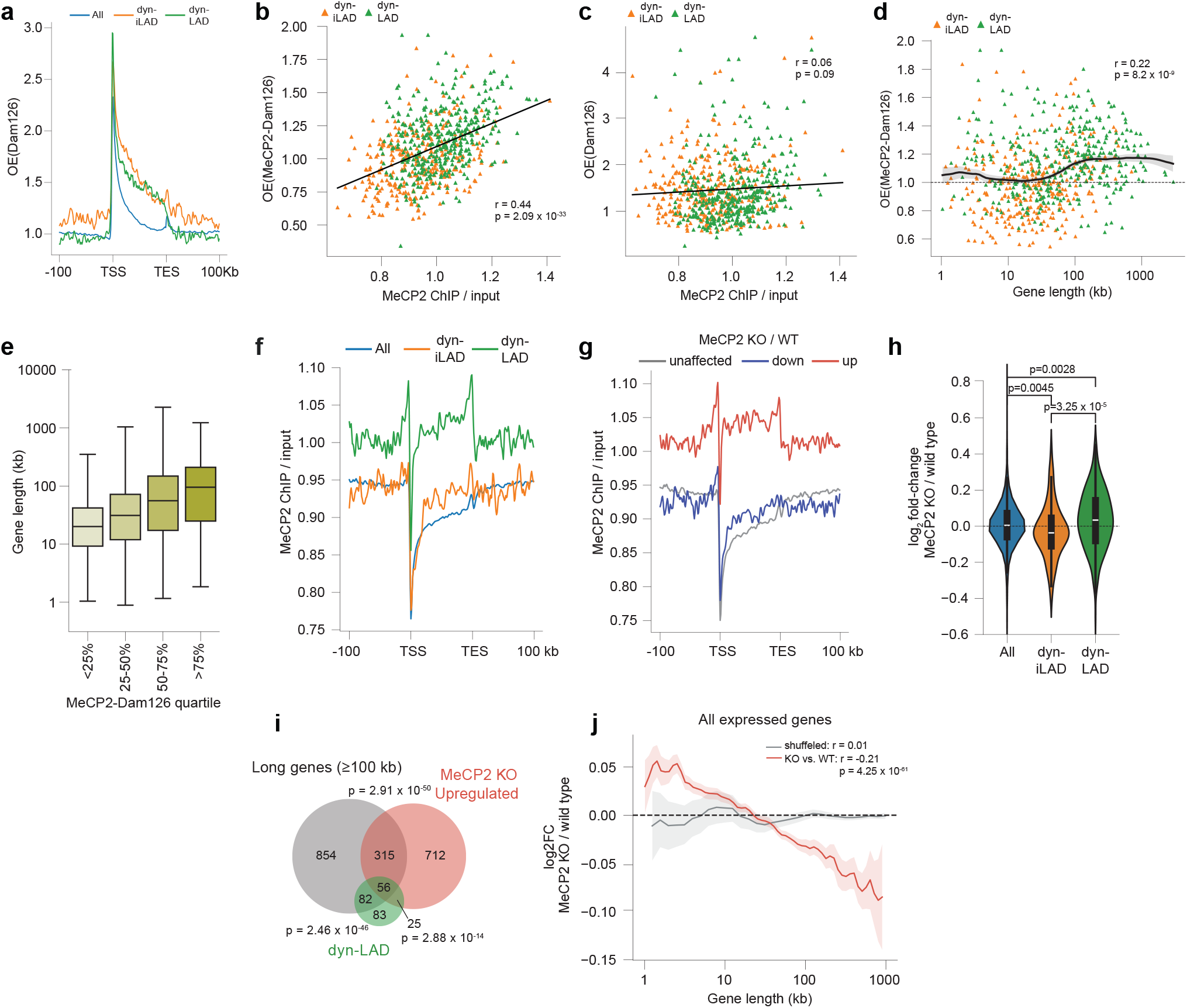
Developmental stage-specific regulation of long nuclear lamina genes by MeCP2. **a**, Dam126 signal aligned over all expressed genes, dyn-iLAD and dyn-LAD genes. **b**,**c**, Correlation between MeCP2 ChIP-seq^46^ and MeCP2-Dam126 (**b**) or Dam126 (**c**) quantified on dyn-iLAD and dyn-LAD genes. Pearson correlation (r) and p-value from null hypothesis test are shown. **d**, MeCP2-Dam126 enrichment on dyn-iLAD and dyn-LAD genes as a function of gene length. Black lines represent the sliding window means with 95% confidence interval in the shaded area. Pearson correlation (r) and p-value from null hypothesis test are shown. **e**, Boxplot of gene length across MeCP2-Dam126 signal enrichment quartiles. **f**,**g**, MeCP2 ChIP-seq of the cortex^46^ aligned over all expressed genes and dyn-iLAD and dyn-LAD genes (**f**) or for genes that are unaffected, downregulated or upregulated upon MeCP2 KO in the cortex^46^ (**g**). **h**, Gene expression changes assessed by RNA-seq for all expressed genes, dyn-iLAD and dyn-LAD genes of the maturing cortex from MeCP2 KO (n=10) and WT littermate control mice (n=10) that are 8 weeks of age^46^. P-values are from a two-sided t-test. **i**, Venn diagram of long genes, MeCP2 KO upregulated genes and the dyn-LAD genes. P-values from hypergeometric test. **j**, Expression changes for all expressed genes as a function of gene length assessed by microarray in the cortex of E15.5 MeCP2 KO (n = 3) and WT control (n = 3) embryos^44^. Red (KO vs. WT) and black (shuffled) lines represent the sliding window means with 95% confidence interval in the shaded area. Pearson correlation (r) and p-value from a null hypothesis test are shown.

**Extended Data Figure 8.**
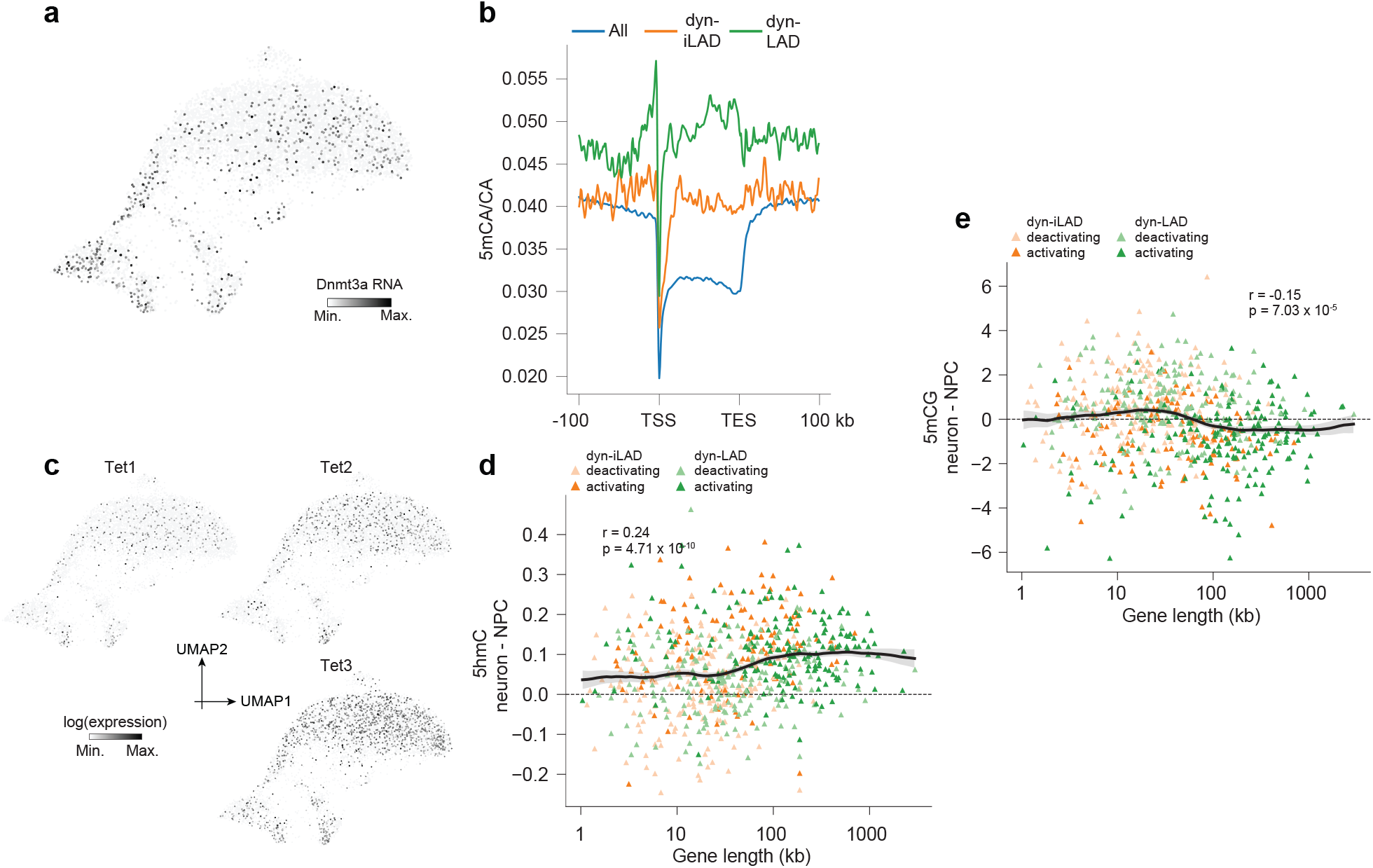
Developmental stage-specific DNA methylation of long nuclear lamina regulated genes. **a**, UMAP plot colored by *Dnmt3a* gene expression. **b**, 5mCA signal quantified by BS-seq^46^ aligned over all expressed genes, dyn-iLAD and dyn-LAD genes. **c**, UMAP plot colored by TET protein gene expression. **d**,**e**, The difference of 5hmC (**d**) or 5mCG (**e**) levels between neurons and NPCs of E15.5 mouse embryos measured by hMeDIP^36^ or Methyl-HiC^80^ on dyn-iLAD and dyn-LAD genes as a function of gene length. Black lines represent the sliding window means with 95% confidence interval in the shaded area. Pearson correlation (r) and p-value from a null hypothesis test are shown.

## Methods

### Animal experiments

All animal experiments were approved by the Dutch Animal Experiments Committee (Dier Experimenten Commissie, DEC), performed in line with institutional guidelines of University Utrecht and UMC (University Medical Center) Utrecht and conducted in agreement with Dutch law (Wet op de Dierproeven, 1996) and European regulations (Directive 2010/63/EU). C57BL/6J mice were obtained from the Jackson laboratories (strain 000664), bred in-house and used for all experiments. All mice were raised with their mothers up to 4 weeks of age, maintained in 12 hour light-dark cycles at 22 ± 1 °C and fed without restriction (standard laboratory chow, Special Diet Services (SDS), CRM (E)). Mice were housed with companions in transparent plexiglas cages with wood-chip bedding and paper tissue for nest building and cage enrichment. Embryonic day 0.5 was established as the day of the vaginal plug. Both female and male embryos were used throughout the study.

### Plasmids

During in utero electroporation (IUE), we co-injected DamID constructs with a pEGFP-N1 vector (Clontech), to assess the expression of the exogenous constructs during tissue dissections and FACS. To express the Dam-Lamin B1 fusion protein, we custom designed a construct mRuby2-P2A-HA-miniAID-Dam-V5-LaminB1 construct (Genewiz), followed by a SV40 poly-A signal, which was cloned under an EF1α promoter sequence. To create this construct, an in-house pCCL-sin-EF1α-Dam-LaminB1 vector was used, in which the original Dam-Lamin B1 sequence was replaced by mRuby2-P2A-HA-miniAID-Dam-LaminB1, while other sequences in the vector (WPRE, SV40 poly-A, backbone) were kept intact. To express the untethered Dam, an in-house version of the Ai9 plasmid (Addgene, #22799) was used, placing the mRuby-P2A-HA-miniAID-Dam126 construct under the CAG promoter. In this vector, a β-globin polyadenylation signal is included after the Dam126 sequence. The Dam126 mutant sequence^20^ originated from a vector from^57^. For the CAG-MeCP2-Dam126 construct, the wildtype murine *Mecp2* gene sequence was generated from the pEGFP-N1_MeCP2(WT) plasmid (Addgene, #110186) through PCR amplification (primers fw, aaaatcgatttgaggatgtataaagaactagcgctaccggact; rev atttttcttcatggtgctgcgatcgaagaccggtggatccttacagctaactct). This resulted in a 1551 bp fragment, which was cloned in the CAG-mRuby2-P2A-HA-mAID-Dam126 vector at the 5’ end of the Dam126 sequence through Gibson cloning, to generate CAG-mRuby2-P2A-HA-mAID-MeCP2-Dam126. Sequences of all vectors were verified by Sanger sequencing (Macrogen).

### In utero electroporation of embryonic cortex

In utero electroporation (IUE) was performed as previously described^69^. Briefly, E13.5, E14.5 or E16.5 pregnant C57BL/6J mice were put under deep anesthesia using Isoflurane (Zoetis, induction - 3-4% in sleeping compartment; surgery - 1.5-2.0% in mouth mask; always 30% O2) and injected with 0.05 mg/kg buprenorfinhydrochloride (in saline) 30 minutes before the operation. Under sterile conditions, the abdominal cavity was opened to expose the uterine horns. For the Dam-Lamin B1 injections, a DNA solution was prepared containing the EF1α-mRuby2-P2A-HA-miniAID-Dam-LaminB1 construct at either 0.5 μg/μl or 2.0 μg/μl and the GFP construct at 0.4 μg/μl. mRuby expression for each embryo was independently evaluated at FACS, before deciding with the optimal concentration to proceed. For the Dam126 injections, a DNA solution was prepared containing the CAG-mRuby2-P2A-HA-miniAID-Dam126 construct at 0.5 μg/μl, in combination with the GFP construct at a 0.25 μg/μl. For the MeCP2-Dam126 injections, a DNA solution was prepared containing the CAG-mRuby2-P2A-HA-miniAID-MeCP2-Dam126 construct at either 1.0 or 0.5 μg/μl, in combination with the GFP construct at a 0.25 μg/μl. For all conditions, the DNA constructs were dissolved in MiliQ water with 0.05% Fast Green (Sigma, F7252) and were injected at a volume of 1.7 μl unilaterally into the lateral ventricle of embryos using glass micro-pipettes (Harvard Apparatus) and a PLI-100 Picoinjector (Harvard Apparatus, Holliston, MA). The brain (motor cortex) was then electroporated with gold-plated tweezer electrodes (Fischer Scientific) and an ECM 830 Electro-Square-Porator (Harvard Apparatus, Holliston, MA) set to 5 unipolar pulses of 50 ms pulse length at 30 V (950 ms interval). Next, embryos were placed back into the abdomen, and abdominal muscles and skin were sutured separately. The whole procedure from anesthesia to awaking of the mother by release from Isoflurane took approximately 1 hour. The mice were monitored daily up until 48 hours after IUE (at E15.5, E16.5 or E18.5), when the mothers were sacrificed through cervical dislocation.

### Cryosectioning

Mothers were sacrificed at E16.5 (48 hours after IUE) and whole brains from the pups were collected for immunostainings in ice-cold Leibovitz’s L-15 medium (Gibco, 11415064), washed in PBS and fixed by immersion in 4% paraformaldehyde (PFA, Sigma, 30525-89-4) in PBS at 4 °C overnight. Brains were then washed in PBS, cryoprotected in 30% sucrose in PBS at 4 °C and frozen in 2-methylbutane (Sigma, 277258d), after which they were embedded in Embedding matrix (Thermo Fisher Scientific). Brains were sliced into 25 µm coronal sections using a cryostat (Leica, CM1950 Ag Protect) set at −14 °C, mounted on slides (VWR Superfrost Plus, 631-0108), air-dried and stored at −80 °C.

### Immunostaining and microscopy

Cryosection slides were hydrated in PBS with 0.5% TritonX-100 (PBS-T, Sigma, X100-100ML) for 10 min at room temperature (RT) after which excess PBS-T was removed. Slides were treated with 400 μl blocking buffer (2% freeze-dried normal donkey serum (NDS, Jackson Immunoresearch, 017-000-121) in PBS-T) at RT for 30 min in a humidifying chamber followed by primary antibody incubation in 250 μl of blocking buffer at RT for 2 hours in a humidifying chamber. Primary antibody used is Rabbit anti-HA-Tag (Cell Signaling Technology, C29F4) at 1 in 500 dilution. Slides were washed three times with PBS before secondary antibody incubation in 300 μl blocking buffer at RT overnight in a humidifying chamber. Secondary antibody (at 1 in 1000 dilution) used is Donkey anti-Rabbit Alexa647 (Invitrogen, A-31573). Sections were washed three times with PBS, counterstained with 1 µg/ml DAPI and mounted in ProLong Glass Antifade mounting medium (Thermo Fisher Scientific, P36980). Images were acquired using a Leica TCS SPE confocal microscope using the standard Leica LAS-X software and were processed with ImageJ (version 2.14.0).

### RNA FISH

E16 embryos (n=3) were fixed, cryoprotected and sectioned at 30 μm thickness as indicated above. Sections were preserved at −70 °C, air dried at room temperature for up to one hour prior to the procedure. In situ hybridization was performed using the viewRNA tissue kit (Thermo Fisher Scientific, #QVT4700), following the manufacturer’s instructions with minor modifications. Specifically, the embryonic slides were prefixed with 4% PFA for 15 min and washed in PBS for 3 x 5 min prior to the process; The recommended 400 µl per slide was reduced to a volume of 200 µl per slide for longevity; target probes were hybridized overnight. Hybridization steps were conducted in the HybEX II Oven (ACD, Bio-Techne) at 40 °C. Sections were washed 3 x 5 min in PBS and blocked in 400 µl of blocking buffer (2% NDS in PBS with 0.1% TritonX-100) for 30 min. Sections were incubated with 0.5 µg/ml rabbit anti-Lamin B1 antibody (Abcam, #ab16048) in blocking buffer at RT for 2 hours, washed 3 x 30 min in PBS and stained with 0.5 µg/ml donkey anti-rabbit-Alexa488 antibody (Life Technologies, #A21206l) for 2 hours. Sections were washed three times with PBS, counterstained with 1 µg/ml DAPI and mounted in Fluorsave (Sigma-Aldrich, #345789) mounting medium. Intron-specific probes for *Hopx, Nrp1, Vim, Satb2, Pax6* and *Nav3* were designed against selected 5 kb – 15 kb intronic sequences (Supplementary Table 1), generated by the provider (Thermo Fisher Scientific). Regions were selected by presence in all known isoforms and evolutionary conservation.

Imaging experiments were performed using a Nikon TI inverted microscope with NIS Element Software equipped with a perfect focus system, a Yokagawa CSU-X1 spinning disc, an iXon Ultra 897 EM-CCD camera (Andor), and a motorized piezo stage (Nanocan SP400, Prior). To create large overview images, a grid of 90% overlapping individual field of views (FOVs) in a single z-plane were acquired with a 20x 0.75 NA air objective and stitched together using the NIS software. High-resolution images for distance quantifications were acquired using a 100x 1.49 NA oil-immersion objective and 15-21 z-planes with 0.5 μm spacing, depending on the thickness of the section.

### Embryonic cortex dissociation and single-cell collection for scDam&T-seq

Mothers were sacrificed 42-48 hours after IUE (at E15.5, E16.5 or E18.5) and whole brains from the pups were collected in Leibovitz’s L-15 medium. Cortices were dissected under a stereo microscope (Leica) and dissociated using the Papain Dissociation System (Worthington Biochem, LK003150) with 25 µg/ml DNaseI (Worthington) for 10 min at 37 °C, followed by mechanical dissociation and another incubation at 37 °C for 5 min, before final dissociation into single cells. Cells were washed in 10 ml of Leibovitz’s L-15 medium supplemented with 2% fetal bovine serum (FBS, Sigma, F7524) to stop enzymatic digestion. Cells were centrifuged at 200 g for 7 min at 4 °C, resuspended in Leibovitz’s L-15 medium and pipetted through a Cell Strainer Snap Cap into a Falcon 5 ml Round Bottom Polypropylene Test Tube (Fisher Scientific, 10314791). Cells were kept on ice from this point onwards. DAPI (Sigma, D9542) was added to a final concentration of 0.5 µg/ml just prior to FACS sorting with the BD Influx Cell Sorter. GFP and mRuby double-positive cells were sorted as single cells in 384-well PCR plates (BioRad, HSP3831) containing 50 nl CEL-seq2 primer (1500 nM, as described in^17^) and 5 µl mineral oil (Sigma, M8410) per well. After sorting, plates were sealed with aluminum covers (Greiner, 676090), spun for 2 minutes at 2000 g at 4 ^°^C and stored at −80 °C.

### scDam&T-seq processing

Uniquely-barcoded CEL-seq2 primer and DamID adapter sequences (top and bottom oligos) are described in^18^. The scDam&T-seq processing was largely performed as previously described in^17^, but reaction volumes of all mixes were equimolarly halved (0.5x) to reduce overall processing costs. As such, the added volumes per well were 50 nl CEL-seq2 primer (before sorting, with 5 µl mineral oil), 50 nl lysis mix, 75 nl reverse transcriptase mix, 925 nl second strand mix, 250 nl proteinase K mix, 150 nl DpnI mix, 50 nl of DamID adapter (1 µM concentration, 25 nM final during ligation) and 450 nl scDam&T T4 ligase mix. The overall composition of the mixes remains unaltered and the incubation times and temperatures are as described. Briefly, cells were lysed and reverse transcription was performed to generate cDNA-mRNA hybrids, by hybridizing the poly-A tail of the mRNA to the poly-T sequence in the cell-specifically barcoded CEL-seq2 primers. This was followed by second-strand synthesis to generate a double stranded and barcoded cDNA. After a proteinase K step to remove all chromatin proteins and generate ‘naked’ genomic DNA, methylated adenines (GATC) resulting from Dam enzyme activity were specifically digested with DpnI and double-stranded DamID adapters containing cell-specific barcodes were ligated to these genomic fragments.

### Library preparation

After scDam&T-seq plate processing, all cells with non-overlapping barcodes (generally a full 384-well plate) were pooled for in vitro transcription (IVT), generally following the protocol as previously described in^17^. To pool cells, plates were spun upside down in a clean VBLOK200 Reservoir (ClickBio) for 4 minutes at 300 g, before collecting the complete water phase with residual mineral oil in one tube. Residual mineral oil was removed by spinning the sample for 2 minutes at 2000 g and transferring the liquid phase to a clean tube, which was repeated three times. After pooling, samples were incubated for 10-20 minutes with 0.8 volume CleanNGS magnetic beads (CleanNA, CPCR-0050), diluted 1:10 in bead binding buffer (20% PEG 8000, 2.5 M NaCl, 10 mM Tris–HCl, 1 mM EDTA, 0.05% Tween 20, pH 8.0 at 25 °C)). Samples were placed on a magnetic rack (DynaMag™-2, Thermo Fisher Scientific, 12321D) to wash beads two times with 80% ethanol (Boom, 84028185) before allowing beads to dry before resuspending in 8 µl water. IVT was performed by adding 12 µl IVT mix from the MEGAScript T7 kit (Invitrogen, AM1334) for 14 hours at 37 ^°^C, cooling to 4 ^°^C and transferred to ice. Samples were first eluted from the beads and subsequently cleaned with 0.8 volume CleanNGS magnetic beads (undiluted) and three 80% ethanol washes, before eluting in 22 µl water and placing samples on ice. Next, the RNA was fragmented by adding 5.5 µlfragmentation buffer (200 mM Tris-acetate pH 8.1, 500 mM potassium acetate, 150 mM magnesium acetate) and incubating samples at 94 ^°^C for 105 seconds before immediately transferring to ice. The reaction was stopped by adding 2.75 µl STOP buffer (0.5 M EDTA pH 8). After transferring samples back to room temperature, samples were cleaned with 0.8 volume CleanNGS magnetic beads (undiluted) and three 80% ethanol washes, before eluting in 8.5 µl water – the clean aRNA sample. aRNA quality is evaluated on a Bioanalyzer RNA 6000 Pico Kit chip (Agilent, 5067-1513) and concentrations were measured using a NanoDrop™ 2000. Library preparation was subsequently performed as described previously^17^, using 5 µl of aRNA and 8 to 10 PCR cycles, depending on aRNA yield. After PCR, samples are cleaned twice with 0.8 volume CleanNGS magnetic beads (undiluted) and finally eluted in 8-12 µl DNAse-free water. Library quality is evaluated on a Bioanalyzer High Sensitivity DNA Analysis chip (Agilent, 5067-4626) and concentrations were measured using a Qubit dsDNA Quantification Assay (Invitrogen, Q32851) on a Qubit 3.0 Fluorometer (Invitrogen). Libraries were run on an Illumina NextSeq500 platform (high output 2 x 75 bp) or an Illumina NextSeq2000 platform (high output 2 x 100 bp), at an intended output of 1.0 - 1.5 × 10^5^ read-pairs per cell.

### Processing of scDamID and scDam&T-seq data

Data generated by the scDam&T-seq protocol was largely processed following the workflow and scripts described in^17^. Scripts and a workflow description are also available on www.github.com/KindLab/scDamAndTools. The procedure is described in short below.

### Demultiplexing

All reads are demultiplexed based on the barcode present at the start of R1 using a reference list of barcodes. The reference barcodes contain both DamID-specific and CEL-seq2-specific barcodes and zero mismatches between the observed barcode and reference are allowed. The UMI information, also present at the start of R1, is appended to the read name.

### DamID data processing

DamID reads are aligned using bowtie2 (v. 2.3.3.1)^70^ with the following parameters: ‘‘--seed 42 -- very-sensitive -N 1.’’ The mm10 reference genome is used for alignment. The resulting alignments are then converted to UMI-unique GATC counts by matching each alignment to known strand-specific GATC positions in the reference genome. Any reads that do not align to a known GATC position or have a mapping quality smaller than 10 are removed. Up to 2 unique UMIs are allowed for single-cell samples to account for the number of alleles. Finally, counts are binned at the desired resolution, generally 100 kb.

### CEL-seq2 data processing

CEL-seq2 reads are aligned using tophat2 (v. 2.1.1)^71^ with the following parameters: ‘‘--segment-length 22 --read-mismatches 4 --read-edit-dist 4 --min-anchor 6 --min-intron-length 25 --max-intron-length 25000 --no-novel-juncs --no-novel-indels --no-coverage-search --b2-very-sensitive - -b2-N 1 --b2-gbar 200.’’ The mm10 reference genome and the GRCm38 (v. 89) transcript models are used. Alignments are subsequently converted to transcript counts per gene with custom scripts that assign reads to genes similar to HTSeq’s^79^ htseq-count with mode ‘‘intersection_strict.’’

### DamID data filtering and normalization

Single-cell DamID samples were filtered based on a construct-specific depth threshold of 5,00 unique GATC’s for Dam126 and 1,000 for Dam-Lamin B1 and MeCP2-Dam126. After filtering, DamID data was normalized by taking the observed DamID count over the expected GATC distribution (OE), as done previously^18^. To obtain a per gene value for each of the DamID constructs, the log2 OE score in the 100 kb bins where the TSS of that gene resides was used, as done previously^18^.

### Analysis of CEL-seq2 data with Scanpy and Harmony

After pre-processing, transcriptomic data was further processed using Scanpy^72^. Cells with less than 100 UMI’s or more than 10% mitochondrial reads, and genes detected in less than 10 cells were excluded from the analysis. Data was subsequently scaled using `scanpy.pp.normalize_total(target_sum=1e4`) and log-transformed. The top 2000 most variable genes were identified using `scanpy.pp.highly_variable_genes(n_top_genes=2000, flavor=‘seurat’, batch_key=‘limsid`‘). Confounding factors such as the total number UMI’s and percentage of mitochondrial reads were regressed out with `scanpy.pp.regress_out` (). Genes were scaled to the unit variance with scanpy.pp.scale(max_value=10). Dimensionality of the data was initially reduced with principal component analysis using `scanpy.tl.pca(svd_solver=‘arpack’, n_comps=100`). The different experimental batches were harmonized using Harmony^73^ with the following parameters `harmony.harmonize(batch_key=‘limsid’)`. A neighborhood graph was computed with `scanpy.pp.neighbors(n_neighbors=40, n_pcs=20, use_rep=‘X_pca_harmony’)`. Clustering of cells was performed with `scanpy.tl.leiden(resolution=.75)`. Finally, PAGA and UMAP embeddings were computed using `scanpy.tl.paga(), scanpy.pl.paga(plot=False)` and `scanpy.tl.umap(min_dist=.3, random_state=0, init_pos=‘paga’)`.

### RNA velocity with Velocyto and scVelo

To distinguish pre-mRNA (unspliced) from mature mRNA (spliced) molecule reads Velocyto was used with default settings^27^. For RNA velocity analysis, default steps of the scVelo package were used^74^.

### DamID binarization and Contact Frequency and combined single-cell heatmap

If specifically stated, data was binarized by setting a threshold of log2(≥1) based on the distribution of OE values of 100 kb genomic bins^75^. The contact frequency (CF) metric is used throughout the manuscript and is defined by calculating the fraction of samples (single cells) that meet the binarization threshold for a given 100 kb bin^75^.

### Definition of dynamically expressed (i)LAD genes

All dynamically expressed genes over pseudotime were identified using graph_test() of Monocle 3 with default settings^23^, and top dynamically expressed genes were subsetted by setting a threshold on the resulting q-value. We noticed that having a single q-value threshold biases genes heavily towards stem cell (deactivating) genes, likely due to the discrete positioning of stem cells in UMAP space. To balance activating and deactivating genes, a separate q-value threshold was set for both (deactivating: 1e-40, activating: 1e-5). Activating and deactivating genes were distinguished by simply calculating the Pearson correlation coefficient of each gene over pseudotime. The resulting set of dynamically expressed genes were grouped into either nuclear lamina associated (i.e. dyn-LAD) or not (i.e. dyn-iLAD), by checking whether a gene was in a LAD in any of the cell types included in the pseudotime analysis, where LAD means a Dam-Lamin B1 CF of ≥ 0.20 in the given cell type.

### Gene Ontology analysis

Gene Ontology enrichment analysis was performed for Mus Musculus using the Gene Ontology database^76^ for biological processes on the identified sets of dyn-iLAD and dyn-LAD genes. All genes were used as a background reference.

### Pseudotime analysis

Pseudotime inference was performed using Monocle3 v.1.0.0^23^ with default settings.

### Plotting gene dynamics over pseudotime

When plotting cells over pseudotime, a rolling mean is shown together with the confidence interval for the mean. To obtain these measurements, we calculated the mean and standard deviations of the metric on the y-axis for each point on the x-axis using a local linear regression approach where data points are weighted according to an exponential decay, that is, exp(−*d*/τ). Here *d* is the distance between the point at the x-axis where the mean is being determined and the data point, and τ is a ‘decay factor’ (or effective radius). The shadings indicate a 95% confidence interval for the means and are determined by 1.96 times the standard deviations, measured using the same exponentially weighted approach as the means.

### Definition of long genes

Gene length was defined by taking the distance between TSS and TTS for the longest isoform of each mm10 Refseq gene. Long genes were defined as larger than 100 kb.

### Definition neural vs. embryonic stem cell-specific LADs

Migrating neurons (MN) and embryonic stem cell (ESC)^35^ Dam-Lamin B1 data were used to identify neural-specific genome–lamina interactions. For this purpose, the CF differential (ΔCF) was taken between MN and ESC Dam-Lamin B1. Of this differential, regions with ≥0.1 MN signal were called neural-specific and the same for ESC specific signal. Regions smaller than 300 kb were excluded and two adjacent regions interrupted by ≤100 kb were merged.

### RNA FISH analysis

The 100X magnified images of the mouse brain tissue slides were used for downstream analysis of the RNA FISH data. A deep-learning based segmentation algorithm Cellpose (version 2.2.2)^77^ was used on the Lamin B1 IF channel to segment the nuclei. The pre-trained ‘cyto3’ model was used with the following parameters `model.eval(diameter=30, flow_threshold=0.5, cellprop_threshold=0, do_3D=False, stich_threshold=0.25)`. RNA FISH probe foci were detected by iterating through the z-slices as follows. Each z-slice was first convolved using `scipy.ndimage.gaussian_filter(sigma=1)`, followed by deriving an Otsu binary threshold to distinguish signal from background pixels. Resulting masks separated by just one pixel were merged using `scipy.ndimage.binary_dilation()`with default parameters. 2D masks were then stitched into 3D using `cellpose.utils.stitch3D(stitch_threshold=0.15)`. Foci smaller than 5 voxels or larger than 500 voxels were excluded from the analysis. The centroid of the foci was determined using `scipy.ndimage.center_of_mass()`. The majority of the foci overlap nuclear segments and the majority of the nuclei have either 0, 1 or 2 foci, rarely more. The distance of the foci centroids to the nuclear periphery was defined as the 3D euclidean distance to the nearest edge of the nuclear segment that the RNA FISH probe focus overlaps. Foci were excluded from the distance analysis if the nearest edge of the nuclear segment represented the edge of the field of view.

### Modeling the temporal relationship between genome–lamina contacts and transcription

To infer the temporal relationship between genome–lamina contacts and gene expression, we modeled gene-specific Dam-Lamin B1 signals and RNA expression levels in a combined dynamic framework, inspired by RNA velocity^74^. Both signals were initially preprocessed by computing their first-order moments with `scvelo.pp.moments(n_pcs=30, n_neighbors=30, use_rep=‘X_pca_harmony’`). We constructed separate models depending on whether the gene was activating or deactivating. For activating genes Dam-Lamin B1 signal (lamina detachment) was modelled as a linear decay:

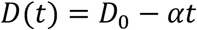

where αis the rate of lamina detachment per unit time *t*. And gene expression was modeled as a delayed exponential increase:

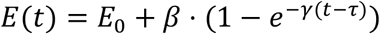

where *β*is the maximum expression change, *γ*the rate of transcriptional activation and τthe delay between lamina detachment and transcription activation.

For deactivating genes, gene expression was modelled as an exponential decay:

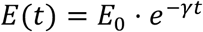

And Dam-Lamin B1 signal (lamina attachment) as as delayed linear increase:

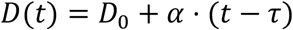

For each gene and model, we implemented a custom cost function to optimize the parameters by minimizing the discrepancy between model predictions and observed data. Full code for the implementation and parameter optimization is available at https://github.com/KindLab/scLADsMouseBrain.

### Quantification along aligned pseudotime

To compare the sequence of gene regulatory dynamics along pseudotime across different datasets (e.g., Dam-Lamin B1 and Dam126 scDam&T-seq), it is important to correct for potential misalignment in pseudotime trajectories for individual genes. Without correction, such misalignment may bias the inferred temporal relationships between different chromatin features measured across datasets with DamID.

To address this, we introduced a metric called aligned pseudotime, which normalizes each gene’s DamID trajectory relative to its jointly measured transcriptional activity in the same cells. We define the moment of transcriptional change as the point where expression reaches the middle 20% of its dynamic range (i.e., between the 40th and 60th percentile), for both deactivating and activating genes. This point is denoted as *t*_()*+_, and we align pseudotime such that *t*_()*+_ = 0.

Formally, for a given gene across pseudotime:

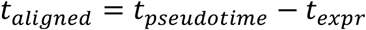

This transformation shifts the observed DamID signal such that dynamics are interpreted relative to transcriptional change. By applying this alignment per gene and per dataset, we can directly compare DamID dynamics across datasets without introducing timing biases due to misaligned pseudotime trajectories.

## Data availability

All genomic and transcriptomic data generated in this study have been deposited at the Gene Expression Omnibus (GEO) under accession number GSE300282.

Previously published RNA-seq of Mecp2 KO mice cortical tissue and MeCP2 ChIP-seq data are available under GSE128186. Previously published MIRA and hMeDIP data are available under GSE38118.

## Code availability

All custom code generated as part of this study have been deposited on GitHub at https://github.com/KindLab/scLADsMouseBrain.

## Acknowledgements

We thank members of the Kind and Basak labs for useful input, discussions throughout the project and critical reading of the manuscript. We thank Nicoletta Landsberger for sharing microarray data. We thank Jeroen J. Pasterkamp for help with IUE design, Nicky Kronenburg for animal caretaking and Solee Pop, Nik Heijmink and Charlotte Valk for experimental assistance. We thank Reinier van der Linden, Anita Pfauth and Joost Korver (Hubrecht FACS facility) for performing single-cell sorting. We thank the Utrecht Sequencing Facility (USEQ) for providing sequencing service and data. USEQ is subsidized by the University Medical Center Utrecht and The Netherlands X-omics Initiative (NWO project 184.034.019). This work was supported by an ERC Consolidator grant (ERC-CoG 1010002885-FateID) to J.K. The Oncode Institute is partially funded by the KWF Dutch Cancer Society. O.B. is supported by the CuCo unusual collaborations grant “Plasticity – here, there, everywhere”, and the NWO Gravitation Grant (BRAINSCAPES, ID: 024.004.012).

## Author contributions

C.M.M. and O.B. conceived combining IUE with scDam&T-seq. P.M.J.R., S.J.A.L, C.M.M., O.B. and J.K conceptualized the project. O.B., K.M.G. and Y.A. performed IUE. S.S.d.V. and C.M.M. performed scDam&T-seq. P.M.J.R., S.J.A.L., S.S.d.V. and O.B. designed RNA FISH experiments. O.B. performed RNA FISH. M.M.M. imaged tissue slides for RNA FISH, assisted by S.S.d.V. P.M.J.R. performed all computational analyses and designed software. E.B. performed preliminary computational analysis of the Dam126 and scATAC-seq data. P.M.J.R., S.J.A.L., S.S.d.V., H.M.B., O.B. and J.K. conceptualized and designed the MeCP2 part of the project. J.G. and M.E.T. supervised H.M.B. and M.M.M., respectively. J.K and O.B. acquired research funding. P.M.J.R. wrote the original manuscript, assisted by S.J.A.L, with input from S.S.d.V., O.B. and J.K.

## Competing interests

The authors declare no competing interest.

## Notes

### Competing Interest Statement

The authors have declared no competing interest.

## References

1. Lodato, S. & Arlotta, P. Generating neuronal diversity in the mammalian cerebral cortex. Annu. Rev. Cell Dev. Biol. 31, 699–720, doi:10.1146/annurev-cellbio-100814-125353 (2015).

2. Di Bella, D. J. et al. Molecular logic of cellular diversification in the mouse cerebral cortex. Nature 595, 554–559, doi:10.1038/s41586-021-03670-5 (2021).

3. La Manno, G. et al. Molecular architecture of the developing mouse brain. Nature 596, 92–96, doi:10.1038/s41586-021-03775-x (2021).

4. Greig, L. C., Woodworth, M. B., Galazo, M. J., Padmanabhan, H. & Macklis, J. D. Molecular logic of neocortical projection neuron specification, development and diversity. Nat. Rev. Neurosci. 14, 755–769, doi:10.1038/nrn3586 (2013).

5. Mannens, C. C. A. et al. Chromatin accessibility during human first-trimester neurodevelopment. Nature, doi:10.1038/s41586-024-07234-1 (2024).

6. Smith, Z. D., Hetzel, S. & Meissner, A. DNA methylation in mammalian development and disease. Nat. Rev. Genet. 26, 7–30, doi:10.1038/s41576-024-00760-8 (2025).

7. Peric-Hupkes, D. et al. Molecular maps of the reorganization of genome-nuclear lamina interactions during differentiation. Mol. Cell 38, 603–613, doi:10.1016/j.molcel.2010.03.016 (2010).

8. Robson, M. I. et al. Tissue-Specific Gene Repositioning by Muscle Nuclear Membrane Proteins Enhances Repression of Critical Developmental Genes during Myogenesis. Mol Cell 62, 834–847, doi:10.1016/j.molcel.2016.04.035 (2016).

9. Poleshko, A. et al. Genome-Nuclear Lamina Interactions Regulate Cardiac Stem Cell Lineage Restriction. Cell 171, 573–587 e514, doi:10.1016/j.cell.2017.09.018 (2017).

10. Ahanger, S. H. et al. Distinct nuclear compartment-associated genome architecture in the developing mammalian brain. Nat. Neurosci. 24, 1235–1242, doi:10.1038/s41593-021-00879-5 (2021).

11. Shah, P. P. et al. An atlas of lamina-associated chromatin across twelve human cell types reveals an intermediate chromatin subtype. Genome Biol. 24, 16, doi:10.1186/s13059-023-02849-5 (2023).

12. Reddy, K. L., Zullo, J. M., Bertolino, E. & Singh, H. Transcriptional repression mediated by repositioning of genes to the nuclear lamina. Nature 452, 243–247, doi:10.1038/nature06727 (2008).

13. Guelen, L. et al. Domain organization of human chromosomes revealed by mapping of nuclear lamina interactions. Nature 453, 948–951, doi:10.1038/nature06947 (2008).

14. Therizols, P. et al. Chromatin decondensation is sufficient to alter nuclear organization in embryonic stem cells. Science 346, 1238–1242 (2014).

15. Leemans, C. et al. Promoter-Intrinsic and Local Chromatin Features Determine Gene Repression in LADs. Cell 177, 852–864 e814, doi:10.1016/j.cell.2019.03.009 (2019).

16. Amir, R. E. et al. Rett syndrome is caused by mutations in X-linked MECP2, encoding methyl-CpG-binding protein 2. Nat. Genet. 23, 185–188 (1999).

17. Markodimitraki, C. M. et al. Simultaneous quantification of protein-DNA interactions and transcriptomes in single cells with scDam&T-seq. Nat. Protoc. 15, 1922–1953, doi:10.1038/s41596-020-0314-8 (2020).

18. Rooijers, K. et al. Simultaneous quantification of protein-DNA contacts and transcriptomes in single cells. Nat. Biotechnol. 37, 766–772, doi:10.1038/s41587-019-0150-y (2019).

19. Aughey, G. N., Estacio Gomez, A., Thomson, J., Yin, H. & Southall, T. D. CATaDa reveals global remodelling of chromatin accessibility during stem cell differentiation in vivo. Elife 7, doi:10.7554/eLife.32341 (2018).

20. Szczesnik, T., Ho, J. W. K. & Sherwood, R. Dam mutants provide improved sensitivity and spatial resolution for profiling transcription factor binding. Epigenetics Chromatin 12, 36, doi:10.1186/s13072-019-0273-x (2019).

21. Taverna, E., Gotz, M. & Huttner, W. B. The cell biology of neurogenesis: toward an understanding of the development and evolution of the neocortex. Annu. Rev. Cell Dev. Biol. 30, 465–502, doi:10.1146/annurev-cellbio-101011-155801 (2014).

22. Buenrostro, J. D. et al. Single-cell chromatin accessibility reveals principles of regulatory variation. Nature 523, 486–490, doi:10.1038/nature14590 (2015).

23. Qiu, X. et al. Reversed graph embedding resolves complex single-cell trajectories. Nat. Methods 14, 979–982, doi:10.1038/nmeth.4402980 (2017).

24. Meuleman, W. et al. Constitutive nuclear lamina-genome interactions are highly conserved and associated with A/T-rich sequence. Genome Res. 23, 270–280, doi:10.1101/gr.141028.112 (2013).

25. Levesque, M. J. & Raj, A. Single-chromosome transcriptional profiling reveals chromosomal gene expression regulation. Nat. Methods 10, 246–248, doi:10.1038/nmeth.2372 (2013).

26. Shah, S. et al. Dynamics and Spatial Genomics of the Nascent Transcriptome by Intron seqFISH. Cell 174, 363–376 e316, doi:10.1016/j.cell.2018.05.035 (2018).

27. La Manno, G. et al. RNA velocity of single cells. Nature 560, 494–498, doi:10.1038/s41586-018-0414-6 (2018).

28. Britanova, O. et al. Satb2 Is a Postmitotic Determinant for Upper-Layer Neuron Specification in the Neocortex. Neuron 57, 378–392, doi:10.1016/j.neuron.2007.12.028 (2008).

29. Jentsch, J. D. & Roth, R. H. The Neuropsychopharmacology of Phencyclidine: From NMDA Receptor Hypofunction to the Dopamine Hypothesis of Schizophrenia. Neuropsychopharmacology 20, 201–225 (1999).

30. Ma, S. et al. Chromatin Potential Identified by Shared Single-Cell Profiling of RNA and Chromatin. Cell 183, 1103–1116 e1120, doi:10.1016/j.cell.2020.09.056 (2020).

31. Gabel, H. W. et al. Disruption of DNA-methylation-dependent long gene repression in Rett syndrome. Nature 522, 89–93, doi:10.1038/nature14319 (2015).

32. King, I. F. et al. Topoisomerases facilitate transcription of long genes linked to autism. Nature 501, 58–62, doi:10.1038/nature12504 (2013).

33. Soheili-Nezhad, S., Ibanez-Sole, O., Izeta, A., Hoeijmakers, J. H. J. & Stoeger, T. Time is ticking faster for long genes in aging. Trends Genet. 40, 299–312, doi:10.1016/j.tig.2024.01.009 (2024).

34. Zylka, M. J., Simon, J. M. & Philpot, B. D. Gene length matters in neurons. Neuron 86, 353–355, doi:10.1016/j.neuron.2015.03.059 (2015).

35. Guerreiro, I. et al. Antagonism between H3K27me3 and genome–lamina association drives atypical spatial genome organization in the totipotent embryo. Nat. Genet. 56, 2228–2237, doi:10.1038/s41588-024-01902-8 (2024).

36. Hahn, M. A. et al. Dynamics of 5-hydroxymethylcytosine and chromatin marks in Mammalian neurogenesis. Cell Rep. 3, 291–300, doi:10.1016/j.celrep.2013.01.011 (2013).

37. Lewis, J. D. et al. Purification, Sequence, and Cellular Localization of a Novel Chromosomal Protein That Binds to Methylated DNA. Cell 69, 905–914 (1992).

38. Sugino, K. et al. Cell-type-specific repression by methyl-CpG-binding protein 2 is biased toward long genes. J. Neurosci. 34, 12877–12883, doi:10.1523/JNEUROSCI.2674-14.2014 (2014).

39. Chahrour, M. et al. MeCP2, a Key Contributor to Neurological Disease, Activates and Represses Transcription. Science 320, 1224–1229 (2008).

40. Mellen, M., Ayata, P., Dewell, S., Kriaucionis, S. & Heintz, N. MeCP2 binds to 5hmC enriched within active genes and accessible chromatin in the nervous system. Cell 151, 1417–1430, doi:10.1016/j.cell.2012.11.022 (2012).

41. Li, Y. et al. Global transcriptional and translational repression in human-embryonic-stem-cell-derived Rett syndrome neurons. Cell Stem Cell 13, 446–458, doi:10.1016/j.stem.2013.09.001 (2013).

42. Skene, P. J. et al. Neuronal MeCP2 is expressed at near histone-octamer levels and globally alters the chromatin state. Mol. Cell 37, 457–468, doi:10.1016/j.molcel.2010.01.030 (2010).

43. Chahrour, M. & Zoghbi, H. Y. The story of Rett syndrome: from clinic to neurobiology. Neuron 56, 422–437, doi:10.1016/j.neuron.2007.10.001 (2007).

44. Bedogni, F. et al. Defects During Mecp2 Null Embryonic Cortex Development Precede the Onset of Overt Neurological Symptoms. Cereb. Cortex 26, 2517–2529, doi:10.1093/cercor/bhv078 (2016).

45. Mellios, N. et al. MeCP2-regulated miRNAs control early human neurogenesis through differential effects on ERK and AKT signaling. Mol. Psychiatry 23, 1051–1065, doi:10.1038/mp.2017.86 (2018).

46. Boxer, L. D. et al. MeCP2 Represses the Rate of Transcriptional Initiation of Highly Methylated Long Genes. Mol. Cell 77, 294–309 e299, doi:10.1016/j.molcel.2019.10.032 (2020).

47. Lister, R. et al. Global epigenomic reconfiguration during mammalian brain development. Science 341, 1237905, doi:10.1126/science.1237905 (2013).

48. Santos, M., Silva-Fernandes, A., Oliveira, P., Sousa, N. & Maciel, P. Evidence for abnormal early development in a mouse model of Rett syndrome. Genes Brain Behav. 6, 277–286, doi:10.1111/j.1601-183X.2006.00258.x (2007).

49. Villard, L. MECP2 mutations in males. J. Med. Genet. 44, 417–423, doi:10.1136/jmg.2007.049452 (2007).

50. Cobolli Gigli, C. et al. Lack of Methyl-CpG Binding Protein 2 (MeCP2) Affects Cell Fate Refinement During Embryonic Cortical Development. Cereb. Cortex 28, 1846–1856, doi:10.1093/cercor/bhx360 (2018).

51. Gulmez Karaca, K., Brito, D. V. C. & Oliveira, A. M. M. MeCP2: A Critical Regulator of Chromatin in Neurodevelopment and Adult Brain Function. Int. J. Mol. Sci. 20, doi:10.3390/ijms20184577 (2019).

52. Baubec, T., Ivanek, R., Lienert, F. & Schubeler, D. Methylation-dependent and - independent genomic targeting principles of the MBD protein family. Cell 153, 480–492, doi:10.1016/j.cell.2013.03.011 (2013).

53. Pantier, R. et al. MeCP2 binds to methylated DNA independently of phase separation and heterochromatin organisation. Nat. Commun. 15, 3880, doi:10.1038/s41467-024-47395-1 (2024).

54. Kriaucionis, S. & Heintz, N. The Nuclear DNA Base 5-Hydroxymethylcytosine Is Present in Purkinje Neurons and the Brain. Science 324, 929–930 (2009).

55. Tahiliani, M. et al. Conversion of 5-Methylcytosine to 5-Hydroxymethylcytosine in Mammalian DNA by MLL Partner TET1. Science 324, 930–935 (2009).

56. Southall, T. D. et al. Cell-type-specific profiling of gene expression and chromatin binding without cell isolation: assaying RNA Pol II occupancy in neural stem cells. Dev. Cell 26, 101–112, doi:10.1016/j.devcel.2013.05.020 (2013).

57. Rang, F. J. et al. Single-cell profiling of transcriptome and histone modifications with EpiDamID. Mol. Cell 82, 1956–1970 e1914, doi:10.1016/j.molcel.2022.03.009 (2022).

58. de Luca, K. L. et al. Genome-wide profiling of DNA repair proteins in single cells. Nat. Commun. 15, 9918, doi:10.1038/s41467-024-54159-4 (2024).

59. Kefalopoulou, S. et al. Time-resolved and multifactorial profiling in single cells resolves the order of heterochromatin formation events during X-chromosome inactivation. bioRxiv, doi:10.1101/2023.12.15.571749 (2023).

60. Ahanger, S. H. et al. Spatial 3D genome organization controls the activity of bivalent chromatin during human neurogenesis. bioRxiv, doi:10.1101/2024.08.01.606248 (2024).

61. van Steensel, B. & Belmont, A. S. Lamina-Associated Domains: Links with Chromosome Architecture, Heterochromatin, and Gene Repression. Cell 169, 780–791, doi:10.1016/j.cell.2017.04.022 (2017).

62. Guarda, A., Bolognese, F., Bonapace, I. M. & Badaracco, G. Interaction between the inner nuclear membrane lamin B receptor and the heterochromatic methyl binding protein, MeCP2. Exp. Cell Res. 315, 1895–1903, doi:10.1016/j.yexcr.2009.01.019 (2009).

63. Fuks, F. et al. The methyl-CpG-binding protein MeCP2 links DNA methylation to histone methylation. J. Biol. Chem. 278, 4035–4040, doi:10.1074/jbc.M210256200 (2003).

64. Kinde, B., Wu, D. Y., Greenberg, M. E. & Gabel, H. W. DNA methylation in the gene body influences MeCP2-mediated gene repression. Proc. Natl. Acad. Sci. USA 113, 15114–15119, doi:10.1073/pnas.1618737114 (2016).

65. Ibrahim, A. et al. MeCP2 is a microsatellite binding protein that protects CA repeats from nucleosome invasion. Science 372, doi:10.1126/science.abd5581 (2021).

66. Liu, S. et al. Cell type–specific 3D-genome organization and transcription regulation in the brain. Sci. Ad. 11 (2025).

67. Dworkin, S. et al. cAMP response element binding protein is required for mouse neural progenitor cell survival and expansion. Stem Cells 27, 1347–1357, doi:10.1002/stem.56 (2009).

68. Goodman, R. H. & Smolik, S. CBP/p300 in cell growth, transformation, and development. Genes Dev. 14, 1533–1577 (2000).

## Methods references

69. van Beuningen, S. F. B. et al. TRIM46 Controls Neuronal Polarity and Axon Specification by Driving the Formation of Parallel Microtubule Arrays. Neuron 88, 1208–1226, doi:10.1016/j.neuron.2015.11.012 (2015).

70. Langmead, B. & Salzberg, S. L. Fast gapped-read alignment with Bowtie 2. Nat. Methods 9, 357–359, doi:10.1038/nmeth.1923 (2012).

71. Kim, D. et al. TopHat2: accurate alignment of transcriptomes in the presence of insertions, deletions and gene fusions. Genome Biol. 14.

72. Wolf, F. A., Angerer, P. & Theis, F. J. SCANPY: large-scale single-cell gene expression data analysis. Genome Biol. 19, 15, doi:10.1186/s13059-017-1382-0 (2018).

73. Korsunsky, I. et al. Fast, sensitive and accurate integration of single-cell data with Harmony. Nat. Methods 16, 1289–1296, doi:10.1038/s41592-019-0619-0 (2019).

74. Bergen, V., Lange, M., Peidli, S., Wolf, F. A. & Theis, F. J. Generalizing RNA velocity to transient cell states through dynamical modeling. Nat. Biotechnol. 38, 1408–1414, doi:10.1038/s41587-020-0591-3 (2020).

75. Kind, J. et al. Genome-wide maps of nuclear lamina interactions in single human cells. Cell 163, 134–147, doi:10.1016/j.cell.2015.08.040 (2015).

76. Thomas, P. D. et al. PANTHER: Making genome-scale phylogenetics accessible to all. Protein Sci. 31, 8–22, doi:10.1002/pro.4218 (2022).

77. Pachitariu, M. & Stringer, C. Cellpose 2.0: how to train your own model. Nat. Methods 19, 1634–1641, doi:10.1038/s41592-022-01663-4 (2022).

78. Li, C. et al. Single-cell brain organoid screening identifies developmental defects in autism. Nature 621, 373–380, doi:10.1038/s41586-023-06473-y (2023).

79. Banerjee-Basu, S. & Packer, A. SFARI Gene: an evolving database for the autism research community. Disease Models & Mechanisms, 133–135 (2010).

80. Noack, F. et al. Multimodal profiling of the transcriptional regulatory landscape of the developing mouse cortex identifies Neurog2 as a key epigenome remodeler. Nat Neurosci 25, 154–167, doi:10.1038/s41593-021-01002-4 (2022).

